# Selection of software and database for metagenomics sequence analysis impacts the outcome of microbial profiling and pathogen detection

**DOI:** 10.1101/2022.04.27.489679

**Authors:** Ruijie Xu, Sreekumari Rajeev, Liliana C. M. Salvador

## Abstract

**Aim:** Shotgun metagenomic sequencing analysis is widely used for microbial profiling of biological specimens and pathogen detection. However, very little is known about the technical biases caused by the choice of analysis software and databases. In this study, we evaluated shotgun metagenomics taxonomical profiling software to characterize the microbial compositions of biological samples collected from wild rodents.

**Method and Results:** Using nine of the most widely used metagenomics software and four different databases, we analyzed shotgun metagenomic sequence data from three sets of wild rodent tissue samples. We demonstrated the discrepancies in results when different databases and software were used, which cause significant variation in microbial community characterizations. Our analysis also showed that these software differed in their ability to detect the presence of *Leptospira*, a major zoonotic pathogen of one health importance.

**Conclusions:** Significant differences in compositional profiles for the same dataset while using different databases and software combinations can result in confounding biological conclusions in microbial profiling.

**Significance and Impact of Study:** This study cautions that the selection of software and databases to analyze metagenomics data can influence the outcome and biological interpretation.

## Introduction

Studies analyzing the composition of microbial communities are frequently used in diverse study fields, such as ecology (Galbraith *et al*., 2018; Grossart *et al*., 2020), agriculture (Mashiane *et al*., 2017; Granjou and Phillips, 2019), human and animal health (Chen *et al*., 2019; Tun *et al*., 2012; Zhong *et al*., 2019), and pharmacology (Chavira *et al*., 2019; Wang *et al*., 2019). Traditional methods used to identify the microbial agents within a biological specimen include methods such as culture (Handelsman, 2004), antigen detection (Desmonts and Remington, 1980; Lequin, 2005), and nucleic acid detection (Yang and Rothman, 2004; Driscoll, 2009) protocols. However, these laboratory methods are limited to studying a single pathogen of interest and lack the ability to scrutinize the community of microorganisms potentially present in a sample. Next-Generation Sequencing (NGS) technologies have provided researchers with a set of culture-independent tools that identify pathogens directly from DNA sequences (Ghosh, Mehta and Khan, 2019) and characterize the diversity and abundance of microbial populations in biological specimens. Hence NGS technologies have emerged as popular tools for microbial profiling and pathogen detection (Tun *et al*., 2012; Skarżyńska *et al*., 2020; Grützke *et al*., 2021).

Taxonomical profiling analysis in the metagenomics discipline uses two popular approaches: the 16S rRNA amplicon sequencing and shotgun metagenomics sequencing (Jovel *et al*., 2016). The 16S rRNA amplicon sequencing method uses polymerase chain reaction (PCR) to amplify hypervariable regions of bacterial 16S rRNA gene and compares these regions to a 16S reference database (DB) (Johnson *et al*., 2019). In contrast, the shotgun metagenomics sequencing-based approach sequences all given DNA present in a sample (Sharpton, 2014). Although lower in cost (Breitwieser, Lu and Salzberg, 2019), 16S rRNA amplicon sequencing is limited to profiling the genomes of bacterial and archaeal taxa, and is subject to amplification biases (Woese, Kandlert and Wheelis, 1990; Janda and Abbott, 2007). On the other hand, the taxonomical profiling using shotgun metagenomics sequence data compares the sequences to reference whole-genome databases. Since the data contain all genetic information present in the sample, this approach avoids the amplification biases observed in 16S rRNA sequencing (Fouhy *et al*., 2016; Ranjan *et al*., 2016) and increases the resolution of microbial identification (Durazzi *et al*., 2021). Most importantly, it has broader applications such as functional profiling and identification of viruses and other microorganisms with simple genomes (Clark and Pazdernik, 2016).

Currently developed shotgun metagenomics sequencing-based taxonomical profiling software can be separated into two groups: the alignment-based and the alignment-free software. Alignment-based software, including BLASTN (Altschul *et al*., 1990; Johnson *et al*., 2008; Camacho *et al*., 2009), which aligns sequences at the nucleotide level, and Diamond (Buchfink, Xie and Huson, 2015), which aligns at the protein level, were thought to have high sensitivity and have been used as the standard for metagenomics profiling. However, these software require a large amount of time and computational resources to build genome alignments for the high number of sequences usually involved in metagenomics profiling studies (Cannings, 2004; Zielezinski *et al*., 2017). Furthermore, recent investigations in alignment-based methods have reported that alignment-based software decrease in sensitivity with the use of mosaic genomes (ex. viruses) (Zielezinski *et al*., 2017). To overcome these limitations, multiple software have been developed using alignment-free algorithms. For example: 1) Kraken2 (Wood, Lu and Langmead, 2019, p. 2) and CLARK (Ounit *et al*., 2015) were designed with k-mer matching algorithms, where only substrings of sequences were matched (Healy and Chambers, 2014); 2) Metaphlan3 (Truong *et al*., 2015; Beghini *et al*., 2021) was designed to identify unique genetic markers within each microbial taxon; and 3) Centrifuge (Kim *et al*., 2016) and Kaiju (Menzel, Ng and Krogh, 2016) were designed to optimize the time and resources of profiling by compressing the reference microbial genomes into the index structures for storing and searching (at the nucleotide and protein levels, respectively) (Burrows and Wheeler, 1994). In addition to the software mentioned above, some methods were developed to improve the results of existing software, such as Bracken (Lu *et al*., 2017) that improves Kraken2’s output by eliminating false positive assignments using a Bayesian framework, and CLARK-s (Ounit and Lonardi, 2016) that improves the sensitivity of CLARK with the use of spaced k-mers. Previous benchmarks on shotgun metagenomic sequencing taxonomical profiling software have evaluated the performances of these software using either *in silico* or *in vitro* datasets (Peabody *et al*., 2015; Escobar-Zepeda *et al*., 2018; Ye *et al*., 2019). However, the performances of these software to analyze the microbial profiling and diagnostic applications in biological specimens have been less studied.

In addition, many of the current alignment-free software, such as Kraken2 and CLARK, require a large number of computational resources for DB building and storage. Some software, such as Kraken2, provide an alternative prebuilt DB for users with inefficient computing resources in addition to the software’ standard DB, which allows the profiling analysis to perform on a machine with RAM as low as 8 GB (full standard Kraken2 DB requires ~30 GB RAM). Kraken2 also provides the option to build customized DBs based on the users’ needs. For example, users can include the genomes of the known host in the customized DB or include a collection of incomplete and draft genomes of the microorganisms of interest in the DB to increase the sensitivity and accuracy of the software’s classification results (Ames *et al*., 2015; Pereira-Marques *et al*., 2019a). The effect of using different DBs to classify microbial profiles and their impact on the downstream microbial characterization and pathogen detection have not been addressed in previous benchmarks. For samples collected from wild animals, the microbiome compositions are unknown and potentially contain taxa that do not have genomes available in the reference DB. These situations can become a potential source of technical errors for accurate detection and profiling a sample’s microbiome.

In this study, we compared the microbial profiles of tissue samples from two species of *Rattus (Rattus Rattus* and *Rattus norvegicus*) using different metagenomic software and DBs. Specifically, we 1) compared the taxonomical profiles classified by four DBs and nine metagenomics profiling software listed above, 2) determined the effect of their differences in the downstream analyses and in the result interpretation; and 3) identified the presence of zoonotic pathogens, *Leptospira*, in the rat kidneys that we collected using each software’ profiling results, and compared the diagnosis of the pathogen with that of the traditional laboratory methods.

## Materials and Methods

### Samples

Tissue samples from kidney (K), spleen (S), and lung (L) were obtained from four rats from two different species, *Rattus Rattus* (R28) and *Rattus norvegicus* (R22, R26, and R27). Rats were captured from the island of Saint Kitts (longitude 17.3434° N and latitude – 62.7559°W) following protocols approved by the Ross University School of Veterinary Medicine (RUSVM) IACUC (approval # 17-01-04). DNA was extracted from samples using DNeasy Blood and Tissue Kits (QIAGEN Scientific Inc., MD, USA), following the manufacturer’s protocol.

### Metagenomic shotgun sequencing

DNA sample quality was assessed via analysis of the DNA purity and integrity with the agarose gel. DNA purity (OD260/OD280) and concentration were measured using the Nanodrop and Qubit 2.0. The library for metagenomic sequences was constructed with 1 μg DNA per sample. Sequencing libraries were generated using NEBNext® Ultra™ DNA Library Prep Kit for Illumina following manufacturer’s instructions. The DNA sample was fragmented (350 bp), end-polished, A-tailed, ligated with Illumina sequencing adaptor and amplified with the PCR technique. The PCR products were then purified for sequencing. Before sequencing, samples were clustered on a cBot Cluster Generation System, then sequenced on an Illumina HiSeq platform for paired-end reads.

### Data pre-processing

Sequencing adapters, low-quality reads, and host DNA reads within the metagenomic samples were removed using the software KneadData (The Huttenhower Lab, no date) with the default Trimmomatic (Bolger, Lohse and Usadel, 2014) (version 0.33) settings (SLIDINGWINDOW:4:20 MINLEN:50) and the “—very-sensitive” Bowtie (Langmead *et al*., 2019) (version 2.3) option. The hosts’ reference sequences, which were used to separate host reads from the microbial reads, were downloaded from the NCBI’s RefSeq DB (Human: GCA_000001405.28_GRCh38.p13; *R. norvegicus*: GCF_015227675.2_mRatBN7.2; *R. Rattus*: GCF_011064425.1_RRattus_CSIRO_v1).

### Metagenomic profiling

#### Software

Nine software (BLASTN, Diamond, Kraken2, Bracken, Centrifuge, CLARK, CLARK-s, Metaphlan3, and Kaiju) were chosen to determine the rats tissues’ metagenomic profiles. All software were used with the default settings according to the instruction manuals provided by the developers.

#### Database building

If the software had pre-built DBs, these were downloaded directly from the software’ homepage (BLASTN, minikraken DB of Kraken2, Centrifuge, and Metaphlan3). Otherwise, DBs were built based on the standard instructions provided by the software’ manual (CLARK, CLARK-s, Diamond, and Kaiju), with the exception of software that had their DBs available online with the contribution of the scientific community. In this case, the DBs were downloaded directly from the online resources (standard DB of Kraken2, maxikraken DB of Kraken2, and Bracken). CLARK-s’ DB was required to be built on top of a CLARK DB of the same composition, but when the DB was built with the genomes of Bacteria, Archaea, Viruses, and Human, the building was suspended by the software with the error message “the number of targets exceeds the limit (16383)”. This limitation was reported to the developer of CLARK-s, but it has not been resolved by the time this manuscript was drafted. We bypassed the limitation by building the DB with Bacteria, Archaea, and Viruses genomes separately, and combining the classifications using each DB at end of the analysis. In addition, Metaphlan3, which identifies the microbial taxon with marker genes, does not have an option to build a customized DB, only the marker DB distributed by the developer could be used for profiling. Detailed information about DB building is available in Table I.

**Table I.** Detailed information with the software and DB assessed.

| Type | Software | Version | Databases | Database Building Resources | Average Runtime/Sample | Runtime Resources | Citations (12/22/2021) | Publication Year |
| --- | --- | --- | --- | --- | --- | --- | --- | --- |
| DNA | Blastn | v 2.12.0 | nt (172 GB) | Pre-built* | ~2 hr | 12 threads, 1.75 GB | 10705 | 2009 |
|  | Kraken2 | v. 2.1.2 | miniKraken V2 (8 GB) | Pre-built* | >1 min | 12 thread, 7.46 GB | 893 | 2019 |
|  |  |  | standard (53 GB) | Downloaded** | ~1 min | 12 thread, 50.54 GB |  |  |
|  |  |  | maxikraken2 (150GB) | Downloaded** | ~3 min | 12 threads, 140.23 GB |  |  |
|  |  |  | custom (60 GB) | 12 threads, 60.28 GB, ~26 hrs | ~1 min | 12 threads, 59.36 GB |  |  |
|  | Bracken | v 2.6.1 | standard (2.7 GB) | Downloaded** | >1 min | 12 thread, 0 MB | 384 | 2017 |
|  | CLARK | v.1.2.6.1 | bacteria (archaea) virus human (168 GB) | 12 threads, 404.48 GB, ~43 hr | ~3 min | 12 threads, 136.83 GB | 457 | 2015 |
|  | CLARK-s | v.1.2.6.2 | bacteria (archaea) + viruses | 12 threads, (261.15 + 30.59) GB, ~37 hr | ~3 min | 12 threads, (289.81 + 72.03) GB | 76 | 2016 |
|  | Centrifuge | v. 1.0.4 | h+p+v+c (33 GB) | pre-built* | ~3 min | 12 threads, 30.62 GB | 659 | 2016 |
| Marker | Metaphlan | v. 3.0.13 | mpa_v30_C HOCOPhlan_201901_marker (2.8 GB) | pre-built* | ~2 min | 12 threads, 3.10 GB | 1292 | 2015 |
| Protein | Diamond | v. 2.0.13 | nr (218 GB) | 12 threads, 7.98 GB, ~2 hr | ~5 hr | 12 threads, 9 GB | 4825 | 2015 |
|  | Kaiju | v.1.8.2 | Refseq (234 GB) | 12 threads, 115.75GB, ~5 hr | ~1 min | 12 threads, 60.27 GB | 839 | 2016 |

\***Pre-built:** the database was pre-built by the developer of the software and was distributed with the software release.
\*\***Downloaded:** the database

For this specific analysis, customized Kraken2 DB was built following the manual’s instructions. All the libraries present in Kraken2’s standard DB (which include NCBI RefSeq’s bacterial, archaeal, viral, human genome, and known vectors (UniVec_Core) libraries) were included in the customized DB, with the addition of the genomes from the two rat species: *R. norvegicus* (GCF_015227675.2_mRatBN7.2) and *R. Rattus* (GCF_011064425.1_R*Rattus*_CSIRO_v1).

### Statistical analysis

Metagenomic profiles provided by each software were loaded into R (R Core Team, 2020) for statistical analysis using the package “phyloseq” (McMurdie and Holmes, 2013). Pairwise significant difference assessments were performed using a Wilcoxon signed-rank test implemented in R’s “rstatix” package (Kassambara, 2021), which is a non-parametric statistical hypothesis test used for comparing repeated measurements on a single sample. Alpha (Shannon, 1948; Simpson, 1949) and beta diversity (Bray and Curtis, 1957) indices (Whittaker, 1960) were used to describe the microbiome compositions within and between samples, respectively, and were calculated with the R package “vegan” (Oksanen *et al*., 2013). The differentially abundant (DA) taxa analyzed between samples collected from two different tissues were determined by the R package “DeSeq2” (Love, Huber and Anders, 2014) using the “Wald” test, which normalizes reads classified under each species taxon with the “poscounts” method. The data visualization for the metagenomics profiles was performed using the R package “ggplot2” (Ginestet, 2011). For all statistical analysis, p-values were adjusted with the Holm-Bonferroni method (Holm, 1979). Results with p-adjusted value (padj) < 0.05 were identified as significant.

## Results

### Computational Resources for DB Setup and Microbial Profiling

Details of the DBs used for each software in this study, as well as the associated computational resources and building time, are available in Table 1 and described in the Supplementary Text 1. Alignment-free software, CLARK and CLARK-s required the most computational resources and time for DB building (Table 1). On the other hand, alignment-based software, BLASTN and Diamond, took the longest time for microbial profiling.

### Differences in Microbial Profiles Classified Using Different DBs and Software

The statistical comparisons between microbial profiles of the rat samples when different DBs were used are available in Table SI.1 and the absolute and relative number of reads classified using different DBs is shown in Figure S1. The DBs minikraken, standard, customized, and maxikraken have classified across samples an average number of reads of 10,755 (SD: 20,651), 19,565 (SD: 26,468), 20,073 (SD: 26,880), and 21,401(SD: 27,043), respectively (Table SI.2). The number of reads classified by each DB under the four highest taxa (Eukaryota, Bacteria, Viruses, and Archaea) are presented in Figure 1a-d. The padj values for all comparisons between DBs are available in Table SI.3. Detailed DB comparisons across all taxa at domain, genus, and species levels are described in Supplementary Text 2.

**Figure 1.**
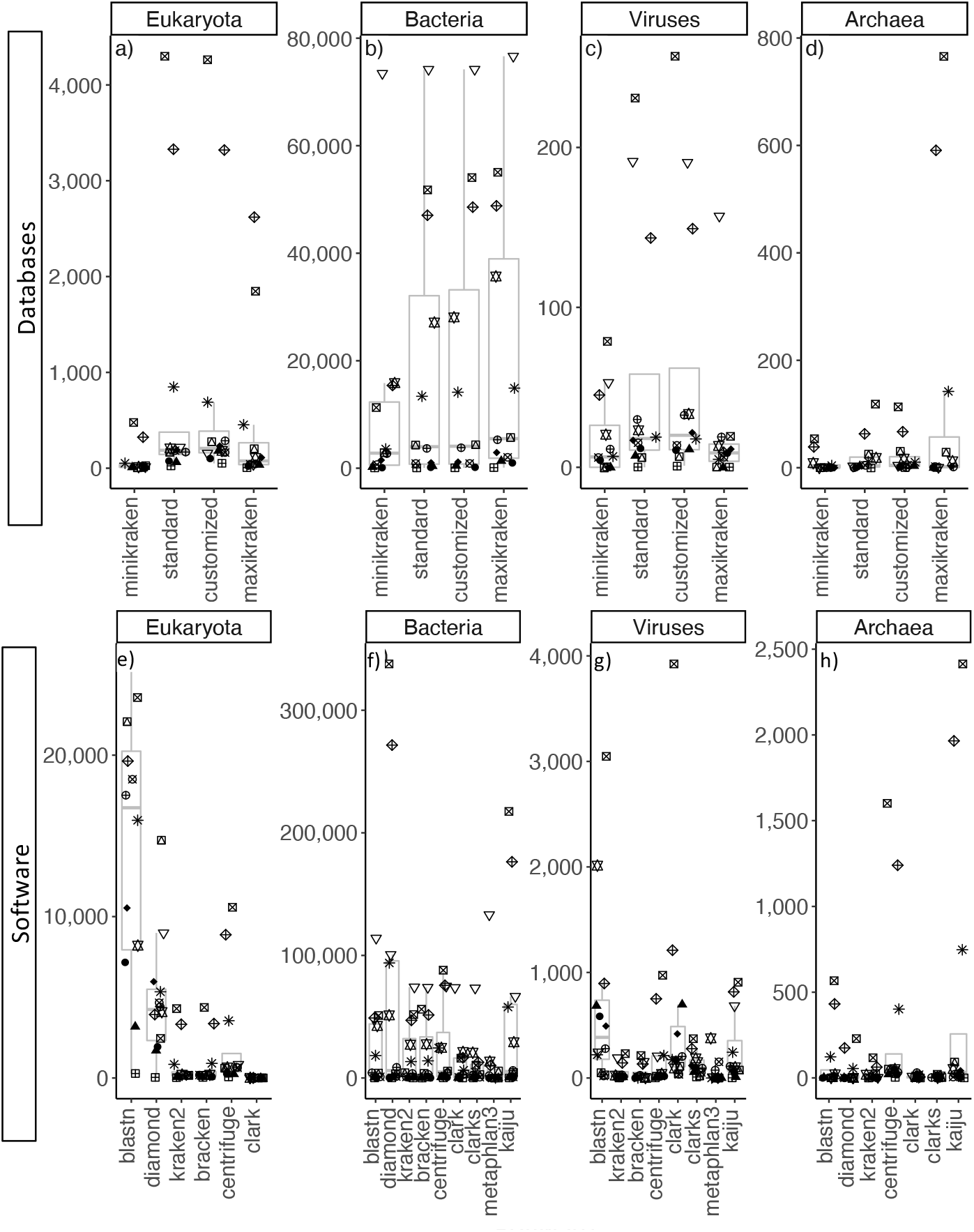
Domain level microbial profiles for rat tissue samples using different DBs (a-d) and software (e-h). All pairwise statistical comparisons between profiles classified by different DBs and software within each domain were performed with a Wilcoxon signed-rank test with p-adj value available in Table SI.3 and Table SII.3 for DBs and software comparison, respectively. Samples: R22.K (●), R26.K (▲), R27.K (◆), R28.K (▽), R22.L 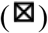, R26.L (✴), R27.L 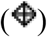, R28.L 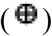, R22.S 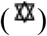, R26.S 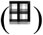, R27.S 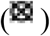, R28.S 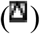.

For comparisons of the microbial profiles classified using different software, the number of total classified reads for each sample at multiple taxonomical levels is available in Table SII.1. The total average number of classified reads ranges from n=10,955 using CLARK-s to n=77,499 using Diamond (Table SII.2). The number of unique taxa classified by each software varied considerably from 18 by Metaphlan3 to 4,816 taxa by Kaiju (Table SII.2). Furthermore, we found that Metaphlan3 did not classify any reads in samples of *Rattus* R26 (R26.K, R26.L. and R26.S) and sample R22.L and R27.K, while other software classified on average 1,252 (SD: 1408), 32,748 (SD: 32,178), 133 (SD: 112), 111,068 (SD: 113,203), and 4,011 (SD: 4,325) reads with these five samples, respectively (Table SII.2).

The number of reads classified by each software at the domain level taxa into Eukaryota, Bacteria, Viruses, and Archaea is shown in Figure 1e-h, and with detailed statistics available in Table SII.3. For the number of reads classified into the Eukaryota taxon, the classifications between all software, except for between Centrifuge and Diamond, showed statistically significant differences. Furthermore, due to the limitation of their DBs compositions, Metaphlan3, CLARK-s, and Kaiju did not report reads classified into the Eukaryota taxon (Figure 1e). For Bacteria taxon classification, the reads classified by CLARK and CLARK-s were found with statistically significant differences in bacterial composition with most other software included in this study (Figure 1f, Table SII.3). Only the classifications by Metaphlan3 and Kaiju were found similar with that classified by CLARK and CLARK’s (Table SII.3). The reads classified into Viruses by different software, on the other hand, were divided into two groups. The first group included the classifications of BLASTN, CLARK, CLARK-s, Metaphlan3, and Kaiju, and the second group included the classifications of Kraken2, Bracken, and Centrifuge. Classification results did not have statistically significant differences within each group, but did between groups (Figure 1g, Table SII.4). Diamond’s classification did not identify any reads as Viruses. Archaea’s read classification was very similar across software (Figure 1h, Table II.3), with the exception of Centrifuge with statistically significant differences from most of other software (BLASTN, Diamond, Kraken2, CLARK, and CLARK-s). Bracken and Metaphlan3 did not classify any reads into the Archaea (Figure 1h).

At the phylum level, the number of unique microbial taxa identified by each software ranged from 5 using Metaphlan3 to 59 using Kaiju. We extracted the top 5 phylum taxa identified from each sample and combined all reads classified to other phyla into the “p__Other_Phyla” taxon (Figure 2). The top 5 Phyla described a large percentage of read classification for all software, but the distribution of reads classified into different phyla taxa are different across software. For example, viral taxon, “p_Pisuviricota”, contributed to over 85% (569/665) of the reads classified in sample R22.K using BLASTN (Figure 2a), but this taxon was not identified by any other software. Metaphlan3 classified all the reads in sample R22.K into “p__Viruses_unclassified” (Figure 2h), and CLARK and CLARK-s classified 63% (120/190) and 57% (95/166) of sample R22.K’s reads into two different viral taxa, “p__Uroviricota” and “p__Artverviricota” (Figure 2f-g). Kaiju also classified 21% of sample R22.K’s reads into “p__Artverviricota” (34/157) (Figure 2i). Similar read distributions involving Virus classification were observed in samples R26.K, R26.S, and R27.K, where BLASTN classified 54% (657/1207), 20% (28/140), and 11% (422/3794) of reads into “p_Pisuviricota” (Figure 2a), respectively; CLARK and CLARK-s classified a large percentage of reads into viral taxon “p__Uroviricota” (CLARK: 71% (636/900), 31/76 (41%), and 18% (201/1099); CLARK-s: 18% (50/271), 18% (7/67), 10% (83/1334), respectively) (Figure 2f-g), but other software only identified zero or a small number of reads into a viral taxon in these samples (Kraken2 classified 4 reads into taxon “p__Uroviricota”, Figure 2c). The distribution of Bacteria reads classified by BLASTN, Kraken2, Bracken, Centrifuge, CLARK, CLARK-s, and Kaiju are relatively consistent across samples. The diversity of taxa identified by Metaphlan3 is significantly lower than the ones identified by other software (Figure 2h). For example, Metaphlan3 identified 100% of sample R27.L’s reads as “p__Proteobacteria” (Genus: *Bordetella*; Species: *B*. *bronchiseptica, B. pertussis, B. parapertussis*, and *B. pseudohinzii*), while other software identified 29% (SD: 12%) of R27.L’s reads as “p__Proteobacteria” on average, from 2-325 unique genus taxa (using Diamond and Kaiju, respectively), and reported *B. bronchiseptica* and *B. pseudohinzii* as the most abundant species under the phylum taxon “p__Proteobacteria”. Diamond’s classification also showed differences in read classification when compared to other software (Figure 2b, Figure S2). The most noticeable difference was the relative abundance of taxon “p__Firmicutes” across samples. In the lung samples, “p__Firmicutes” was classified on average in 17% of R22.L (SD: 9%), 20% of R26.L (SD: 9%), and 14% of R27.L (SD: 8%) by other software from 157 - 971 unique genus and 263 to 2509 unique species, but Diamond only classified 2% (133/4900) of reads as “p__Firmicutes” in sample R26.L (*Lactiplantibacillus plantarum* and *Staphylococcus aureus*), and 0% in both R22.L and R27.L. On the other hand, Diamond identified a relative larger proportion of reads as “p__Firmicutes” in samples R27.S (24%) and R28.L (19%) mostly from the species of genus *Bacillus* when compared to other software (R27.S: mean: 2%, SD: 2%; R28.L: mean: 3%, SD: 3%), which classified most of their Firmicutes reads under the non-*Bacillus* species taxa, except for that of Centrifuge (R27.S: 24%, R28.L: 9%), which also classified most of Firmicutes reads under the species of *Bacillus*.

**Figure 2.**
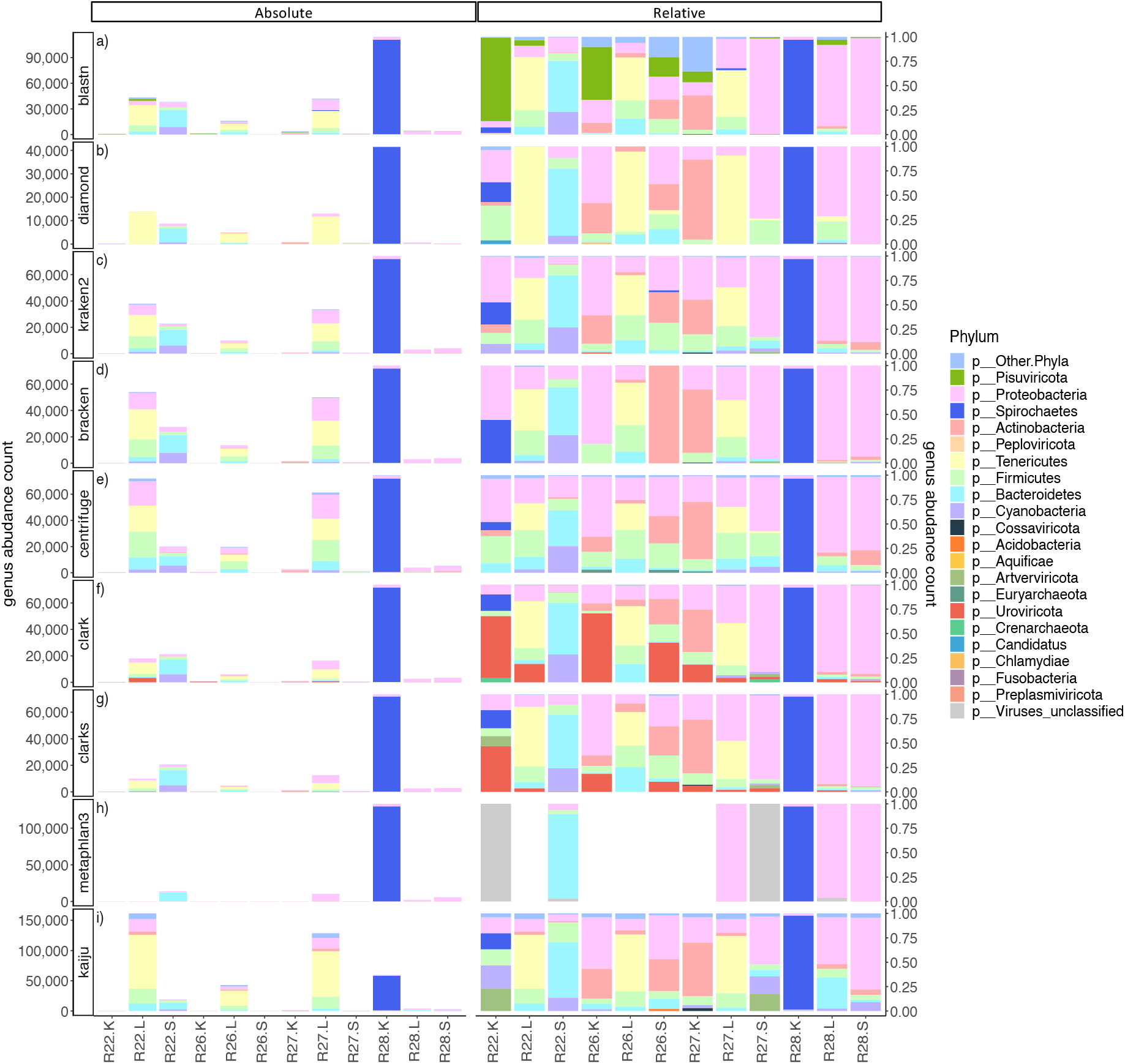
Phylum level microbial profiles for rat tissue samples using nine different software (a-i). Each row panel represent microbial profiles classified using a different software. Left panel is the microbial profiles reported in absolute number of reads and right panel is the microbial profiles reported in relative number of reads. Phylums: P__Other.Phyla 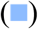, P__Pisuviricota 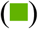, P__Proteobacteria 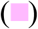, P__Spirochaetes 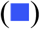, P__Actinobacteria 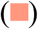, P__Peploviricota 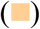, P__Tenericutes 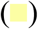, P__Firmicutes 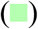, P__Bacteroidetes 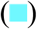, P__Cyanobacteria 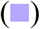, P__Cossaviricota 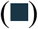, P__Acidobacteria 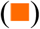, P__Aquificae 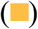, P__Artverviricota 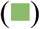, P__Euryarchaeota 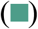, P__Uroviricota 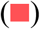, P__Crenarchaeota 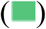, P__Candidatus 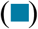, P__Chlamydiae 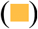, P__Fusobacteria 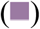, P__Preplasmiviricota 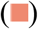, P__Viruses_unclassified 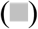.

**Figure 3.**
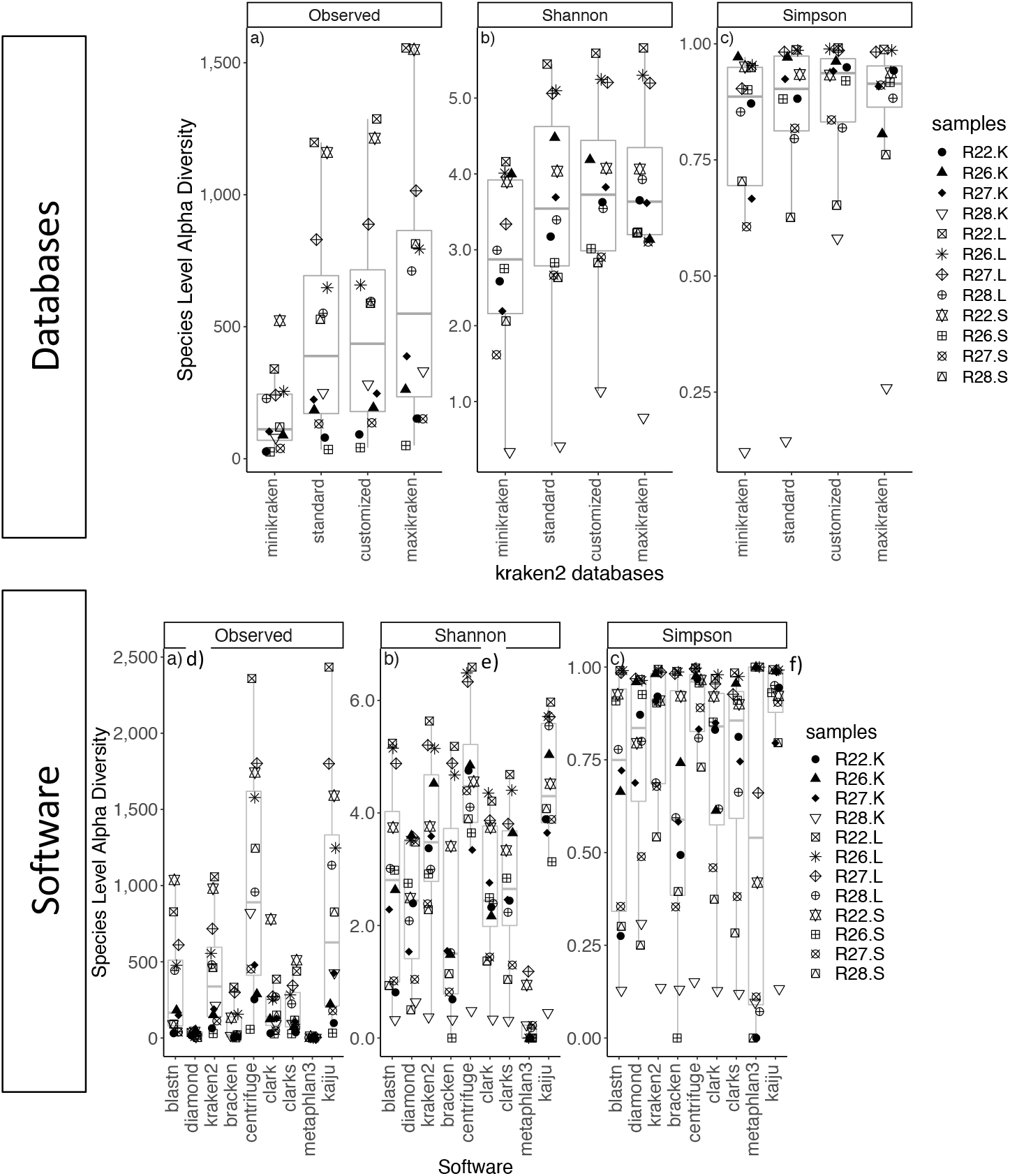
The alpha diversity of the rat samples characterized by different DBs (a-c) and softwares (d-f) is described by the Observed, Shannon, and Simpson indices, which characterize each sample’s microbial composition based on diversity richness and abundance. All three indices were calculated based on the absolute number of microbial reads at the species level. All pairwise statistical comparisons between DBs and software within this figure were performed with a Wilcoxon signed-rank test with p-adj value available in Table SI.4 and Table SII.5 for DBs and software comparison, respectively. Samples: R22.K (●), R26.K (▲), R27.K (◆), R28.K (▽), R22.L 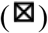 R26.L (✴), R27.L 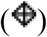, R28.L 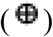, R22.S 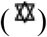, R26.S 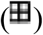, R27.S 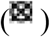, R28.S 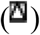.

For the species level classification, the number of reads classified under the same species by each software is available in Table SII.1. Out of all software, Metaphlan3 classified the least number of species taxa, with only 18 species (Table SII.4) while Kaiju classified the most, 4128 species (Table SII.4). From the species level classifications, 9 species taxa were identified by all nine software (*Leptospira interrogans*, *Leptospira borgpetersenii*, *Faecalibacterium prausnitzii*, *Bordetella pseudohinzii*, *Bordetella bronchiseptica*, *Bordetella pertussis*, *Bacteroides uniformis*, *Phocaeicola vulgatus*, and *Bartonella elizabethae*) (Table SII.1). Centrifuge vs Kaiju had the largest overlap of number of identified species taxa (2,285), followed by Kraken2 vs Centrifuge (1,737) and kraken2 vs. Kaiju (1,723) (Table SII.4). The species-level classification of these software mentioned above shared a total of 1,379 species taxa. In addition, BLASTN shared 1,253 species taxa with Centrifuge, 1,207 with Kaiju, and 1126 with Kraken2. CLARK and CLARK-s’ classification shared 1,219 and 1,059 species taxa with Kaiju, respectively. To assess if different software had identified the same species taxa as the most abundant taxa, species taxa with at least 10% of the reads from each sample were selected from each software’ classification. Metaphlan3 identified most of the number of unique species taxa (18), while BLASTN and Kaiju identified the least (7). CLARK vs. CLARK-s and Kraken vs. Bracken shared most of the number of taxa in this category (9 and 8, respectively). Two species taxa were identified by all software as the top ten percent most abundant species taxa, which were *L. interrogans* and *Bartonella elizabethae* (Table SII.1).

### Downstream analyses for microbial community characterization

#### Within-sample diversity (α-diversity)

For within-sample diversity characterization, the observed unique taxa classification results across all four DBs were significantly different from each other (Figure 3a). For species richness characterization within a community using the Shannon indices, only the indices obtained from minikraken DB were significantly different from the results obtained with the other DBs (Figure 3b, Table SI.4). Moreover, the characterization using the Simpson indices was mostly similar between the results of the four DBs (Figure 3c, Table SI.4). Only the Simpson indices obtained from the results of the standard and customized DBs comparison were significantly different (Figure 3c, Table SI.4).

The number of unique observed taxa (Table SII.5, Figure 3d) across different software were largely divergent from each other. Out of the 36 pairwise comparisons between different software, only 6 comparisons were not significantly different (Table SII.5), which were BLASTN’s observed taxa with Kraken2, CLARK, and CLARK-s, comparison between CLARK and CLARK-s, and comparison between Centrifuge and Kaiju. The Shannon indices showed more similarity between software than the number of unique observed taxa, however, they still had 23 out of 36 comparisons between software significantly different from each other (Table SII.5, Figure 3e). The Simpson indices were least impacted by the differences in classification results across the software. Only 7 out of 36 comparisons were found to be significantly different (Table SII.5, Figure 3f). Most of these were identified in comparisons between CLARK-s (3/7) and Centrifuge (4/7) with other software. The Simpson index between CLARK-s and Centrifuge’s classifications were also significantly different from each other.

#### Between-sample diversity (β-diversity)

The pairwise relationships between every two *Rattus* samples were determined with the Bray-Curtis (BC) dissimilarity index, and clustered hierarchically. The BC indices were found to be significantly different across all DBs, except for those reported by the results of maxikraken and customized DBs (Table SII.6). The hierarchical clustering analysis also shows dissimilarities across results when using different DBs (Figure 4a). Three kidney samples (R22.K, R26.K, and R27.K) were found to be clustered with one of the spleen samples (R26.S) in all four DBs’ classifications. However, their relationships with another spleen sample (R27.S) changes with the type of DB used. Despite the differences in the more granular hierarchical clusters, the two major clusters describing the general relationships between samples did not change with the use of different DBs. Three lung samples (R22.L, R26.L, and R27.L) always clustered closely together away from the other samples, while all kidney and spleen samples formed a separate cluster with the other lung sample (R28.L).

**Figure 4.**
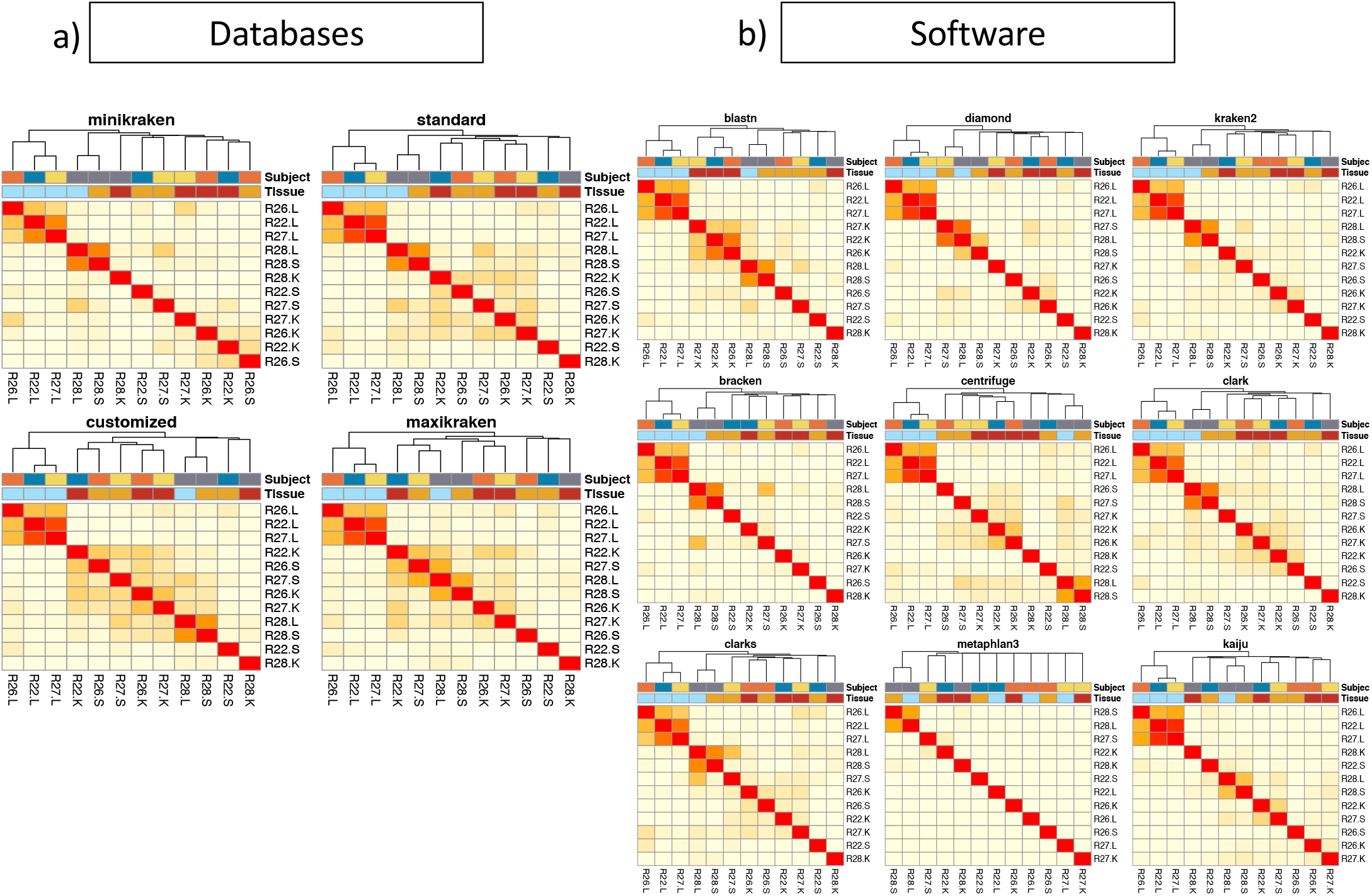
Between-sample microbial composition dissimilarity measure by BC indices using different DBs (a) and software (b). These BC indices were characterized based on the number of microbial reads classified at the species level. Higher BC values indicate a high level of dissimilarity between the two samples’ microbial composition. The red 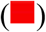 and yellow 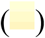 colors show low (0) and high (1) levels of dissimilarity, respectively. Hierarchical clustering was used to cluster together samples that have similar microbial compositions (dendrograms on the left and top of the heatmaps). Pairwise statistical comparisons between DBs and software within this figure were performed with a Wilcoxon signed-rank test with p-adj value available in Table SI.5 and Table SII.6 for DBs and software comparison, respectively.

The hierarchical clusters describing the general relationships between samples remained consistent across all different software (Figure 4b). Except for Metaphlan3, all the other software separated the *Rattus* samples into two large clusters: the first with three lung samples (R22.L, R26.L and R27.L) and the second with a combination of all the kidney and spleen samples, and one lung sample (R28.L). When comparing the BC indices reported by different software (Table SII.6), we found that BLASTN’s BC indices were more similar than the ones of Kraken2, Bracken, and Centrifuge, while the BC indices of CLARK and CLARK-s were more similar with those reported by Diamond, Kaiju, and Kraken2. Metaphlan3, with 5 out of 12 samples unclassified, was significantly different from the other software (Table SII.6).

#### Differentially abundant (DA) taxa identification

DA taxa between samples of different tissues were identified to show the most significantly different microbial taxa between the microbiome of two tissues. For DA taxa identified from lung versus kidney samples at the species level, the number of DA taxon identified by the use of different software ranged from 10 (Diamond) to 596 (Centrifuge) (Table SII.7, Figure 5a). The number of DA taxa identified was higher in in the kidney than in the lung samples for all software’ classifications (Figure 5b). Five significantly abundant species (*Bordetella pseudohinzii, Bordetella bronchiseptica, Leptospira interrogans*, *Leptospira borgpeterseni*, and *Mycoplasm pulmonis*) were classified by all software (Table SII.7). Kaiju and Centrifuge had the highest number of distinct DA taxa (390 and 376 taxa, respectively) (Figure 5a). Although Centrifuge identified the largest number of DA species taxa, Kaiju identified the highest number of unique phylum and genus taxa (42) (Figure 5a).

**Figure 5.**
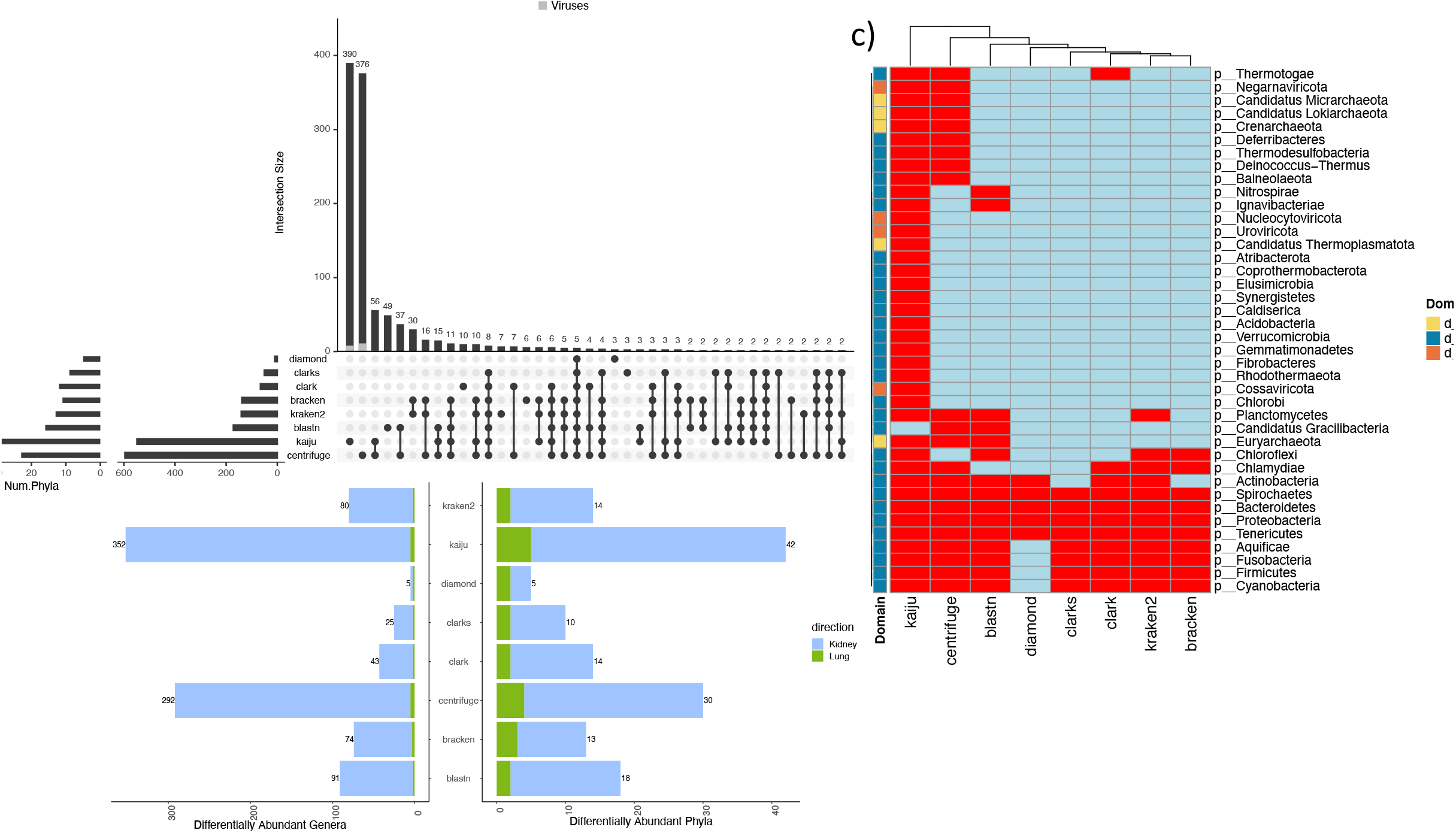
DA microbial taxa identified from comparing the microbial profiles of all rats’ lung and kidney tissues. The number of DA species and phylum identified using different software’s profile is shown at the bar plot directly left to the software names in a). The intersection between DA taxa identified by different software at the species level is shown at the barplot at the top of a), where the number of DA viral taxa were colored in gray 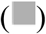. Dotplot at the bottom shown the combinations of intersections between software. Number of DA taxa at Phylum and genus level is shown in b), where numbers of taxa significantly higher in abundance the kidney samples are colored in blue 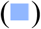, and numbers of taxa higher in the lung samples are colored in green 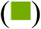. The DA phylum taxa identified by each software is shown in c), where red 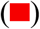indicate the phylum taxa at each row is reported as differentially abundant by the classification of the software in every column, and blue 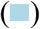 is not reported as DA taxa by the software. Each phylum taxa were also annotated by their corresponding Domain taxa, where dark blue is Bacteria 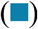, yellow is Archaea 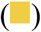, and Orange is Viruses 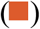.

At the Phylum level lung vs. kidney analysis, the results were consistent for all software except for Diamond, which missed four taxa (“p__Aquificae”, “p__Fusobacteria”, “p__Firmicutes”, and “p__Cyanobacteria”) (Figure 5c). Kaiju and Centrifuge were the only two software that reported viral taxa (“p__Negarnaviricota”, and Kaiju reported “p__Nucleocytoviricota” and “p__Uroviricota”) as DA. Archaeal taxa were only reported by Kaiju, Centrifuge, and BLASTN. All three software reported “p__Euryarchaeota”, and both Kaiju and Centrifuge reported “p__Candidatus Micrarchaeota” and “p__Candidatus Lokiarchaeota”. Finally, Kaiju uniquely reported “p__Candidatus Thermoplasmatota”.

DA taxa were also identified between the microbiomes of lung and spleen (Figure S3) and between kidney and spleen samples (Figure S4). Kaiju and Centrifuge identified most number (484 and 457 species) of DA taxa between lung and spleen samples. The detailed comparisons are described in supplementary Text 3.

### Pathogen detection

The presence of *Leptospira* was identified by all nine software included in the study, however, each software reported *Leptospira* in different samples (Table II). Centrifuge was the only one that reported *Leptospira* in all of the 12 *Rattus* samples, where 9 unique *Leptospira* species were identified (8 from the pathogenic group and 1 from the saprophytic group) (Table SIII.1). Kaiju also identified *Leptospira* from 9 out of 12 samples with 8 unique species (7 from the pathogenic group and 1 from the saprophytic group) (Table SIII.1). Kraken2, following Centrifuge and Kaiju, classified 6 *Leptospira* in 6 samples with 3 unique species all from the pathogenic group (Table SIII.1). Except for Metaphlan3, all software identified *Leptospira* from two of the kidney samples (R22.K and R28.K), which have on average 31 (SD: 3) and 84,344 (SD: 2.2) reads classified under *Leptospira* (Table SIII.2), respectively. BLASTN, Centrifuge, Kaiju, Kraken2 and CLARK identified *Leptospira* from a lung sample (R22.L). Metaphlan3 only identified *Leptospira* in one of the kidney samples (R28.K). All samples identified by at least three software had at least a total of 30 reads classified under *Leptospira* (Table SIII.2). Samples that were only identified by Kaiju or Centrifuge had on average only 2 (R27.K, SD: 1) to 15 (R26.L, SD: 2) reads classified under *Leptospira* (Table SIII.2).

**Table II.**
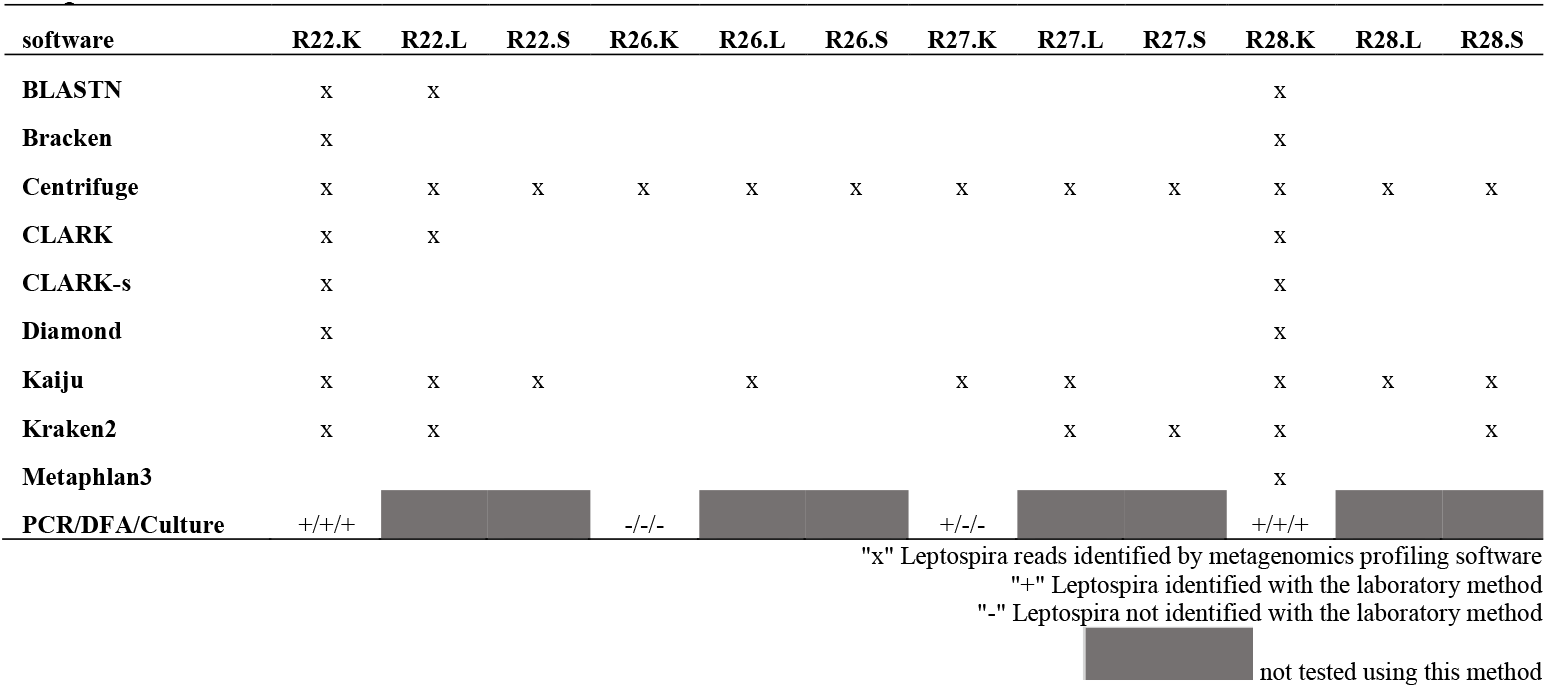
Presence of Leptospira identified from the microbial profiles classified using different software and the presence of Leptospira detected using three traditional methods in the kidney samples.

*Leptospira* detection was also dissimilar when different DBs were used for Kraken2’s classification (Table SIV). Kraken2’s analyses with the maxikraken DB identified *Leptospira* in all samples, while standard and customized DBs identified *Leptospira* in two lung samples (R22.L and R27.L). Standard DB also identified *Leptospira* in the three spleen samples (R22.S, R27.S and R28.S). *Leptospira* detection in the kidney samples using three traditional methods (PCR/DFA/Culture), in comparison to the metagenomics data is shown in Table II.

## Discussion

The field of metagenomics, developed with the advancement of NGS technologies, allows scientists to build a complete and discriminatory microbial profile with viral, archaeal, and bacterial taxa for samples collected from the environments of interest (Jovel *et al*., 2016). These metagenomic profiles can also be used to detect relevant pathogens in clinical and epidemiological investigations (Qin *et al*., 2012; Knights, Lassen and Xavier, 2013) and to observe the interactions between micro-ecosystems and their changing environments (Handley, 2019). Researchers achieve this using a number of bioinformatic analysis software and database combinations. In this study, we identified differences in the microbial profiles when different metagenomics classification software were used. Our results show that there are in fact differences in the classification outputs when different DBs and taxonomical profiling software on the same dataset are used. We conclude that the selection of DBs and methods influences the results of microbiome characterization, and potentially could lead to different biological conclusions and misinterpretation of pathogen presence.

Our study uses biological samples extracted from rats and is different from previous benchmarking studies (Escobar-Zepeda *et al*., 2018; Ye *et al*., 2019), which are based on *in silico* datasets or with the support of laboratory synthetic samples. Our study provides the evidence of reporting false positive or false negative taxonomies if appropriate DBs, or software are not used. Although these differences seem negligible in the benchmarking studies for tools with similar algorithms, they can lead to diverging biological conclusions in the downstream analyses depending on the questions being asked. The biases reported have been understudied; and therefore, it is crucial to demonstrate the effect of these biases with biological data, to raise awareness and identify the potential factors that lead to biased biological conclusions in a metagenomics study.

### Biases introduced by DB selection

Most of the current alignment-free software requires a large number of computational resources for DB building and storage. To allow users with less computational resources to perform the analysis, Kraken2, provides alternative pre-built DBs with only 8 GB. There are also multiple versions of Kraken2’s DBs provided by the science community that can be easily downloaded and updated frequently. For example, the Langmead lab builds the most recent version of Kraken2’s standard DB based on NCBI’s RefSeq library routinely. In addition, the Loman lab has built a Kraken2 DB with the inclusion of draft bacterial, fungal, protozoan, and viral genomes that were not included in the Refseq libraries. Both of these two Kraken2 DBs are freely available online and minimize the workload of building a database from scratch. However, all three DBs mentioned above did not include a reference genome for *Rattus*, which is the host of our dataset. The biases introduced from host genomes included in the DB for metagenomics analysis have been addressed previously (Pereira-Marques *et al*., 2019b). Building a separate DB (customized database for our dataset) with the inclusion of the two *Rattus* hosts genomes on top of the standard DB corrected this issue.

In our analyses with different DBs, although the inclusion of the *Rattus* host genomes did slightly impact the resulting microbial profiles classified from the *Rattus* samples, the discrepancies between the standard DB built with human genome and the customized DB built with *Rattus* genomes were found to be the least among the classification profiles by the four different DBs. The microbial profiles classified using the minikraken and maxikraken DBs are different more significantly from the classifications of standard and customized DBs. The minikraken DB, which is a downsampled DB from the standard DB, classified around only half of the number of reads from the *Rattus* dataset when compared to the classifications of the other three DBs, and were found not capable of classifying some of the most abundant microbial taxa present in the samples. In contrast, the maxikraken DB built with both complete and draft microbial genomes classified the highest number of reads out of the *Rattus* dataset and identified microbial taxa in the samples that were not classified with the other DBs, especially in Bacteria and Archaea’s classifications. Moreover, the maxikraken DB, with the inclusion of other Eukaryota genomes in addition to the human and the *Rattus* genomes, also classified the presence of fungi, parasites, and other Eukaryotic taxa from the *Rattus* dataset. However, both of minikraken and maxiraken DBs were less sensitive in Viruses classifications compared to the standard and customized DBs.

We found that although the number of reads classified using different DBs differed significantly from each other, the characterization of each sample’s microbial communities was not largely biased using different DBs. In our analyses, only the richness of the samples (Shannon indices), which accounts for the rare species within the community, obtained from the minikraken DB, was significantly different from the richness characterized by other DBs. On the other hand, the evenness of each sample’s microbial community measured with the Simpson indices was mostly consistent across all DBs. For microbial communities between samples, we found that only the clusters describing the most distinctive relationships between samples were consistent across the classifications of all DBs. Sophisticated relationships between samples were altered by the biases introduced from DB selection.

### Resources required to use different software

Metagenomics software can be classified into two different categories, alignment-based and alignment-free. The alignment-based software, which suffers greatly from slow speed and the need of large resources, are generally thought to have high sensitivity. On the other hand, the alignment-free software uses relatively small computational resources and significant improvement in speed of the analysis. In our study, the two alignment-based software, BLASTN and Diamond, were the two most time intensive software. They took two and five hours, respectively, on average to complete the analysis for one sample, while other software took at most three minutes for the same task. The time and resources required to build the DBs for the alignment-free software became the trade-off for the speed of the analysis itself. For example, the building of CLARK’s DB took almost two days with 400 GBs of memory used. Fortunately, most of the software included in our study have pre-built DBs distributed with the release of the software (except for CLARK, CLARK-s, Diamond, and Kaiju). However, if the analysis requires the identification of taxa that are not included in these pre-built DBs, the time and resources added to the metagenomics profiling analysis will increase significantly.

### Biases in microbial profiles introduced from software selection

At the Domain level, Eukaryota taxon contributed the most to the dissimilarities between the different software classifications. Almost all pairwise comparisons between the Eukaryota profiles classified by each software were found with no statistically significant differences between each other. Compared to the number of reads classified under Eukaryota, the number of reads classified under Bacteria, Viruses and Archaea by different software were more similar. The classifications of Centrifuge, CLARK, and CLARK-s were frequently identified significantly different from the ones of other software regarding the number of reads mapped to Bacteria and Archaea. The classifications of Viruses, on the other hand, were found separated into two groups where the classifications within a group were similar (group 1: BLASTN, CLARK, CLARK-s, Metaphlan3 and Kaiju; group 2: Kraken2, Bracken and Centrifuge). Diamond did not identify any reads as Viruses. This division in Viruses classifications was further validated at the lower taxonomy levels. The samples with large percentage of reads classified under viral taxa by group 1 software were not profiled by software in group 2. Although software in group 1 were more sensitive in viral identification than of group2 software, the exact viral taxa and the correspondent number of reads using different group1 software were not consistent. The viral taxon identified by BLASTN in high abundance was not identified by any other software included in the analysis. Except for the samples with viral taxa’s classifications, the profiling of Bacteria taxa was found mostly consistent across the software at both Phylum and Genus level. Only the classifications of Metaphlan3, which could only identify a few taxa from each sample with high abundance, and Diamond, which reported conflicting profiles in Firmicutes identification at the Phylum level (*Bacillus* at Genus level) with the classification of all the other software, were different from other software in Bacteria classification.

Compared to Phylum and Genus levels, the classifications at the species level were more divergent across software. Although most software reported more than 1,000 unique species taxa from the *Rattus* profiles (except for Bracken and Metaphlan3), only nine were identified by all software, and only 2 species were found overlapping in taxa with at least 10% in relative abundance.

### Microbial community characterization

In addition to the differences in microbial profiles classified by different software, the differences across the richness of each samples’ microbial community were significant in the majority of the comparisons across software. Most of these were found between the classifications of Kraken2, Metaphlan3, Centrifuge, and Kaiju with other software. However, when richness was measured with species abundance (Simpson index), the characterizations were mostly not affected by use of different software. The characterizations of the relationships between-samples were divergent across software, but the most discriminatory relationships within the rat samples (between the lung and other samples) were captured by most of the software (except for Metaphlan3).

### Differences in differential abundant (DA) taxa

To address potential biases introduced from software selection with biological significance, we identified the DA taxa between samples of different tissues in a pairwise fashion. The classifications of all DA taxa reported at the species level were largely different across software. The largest range in the number of DA taxa reported by different software was found in the analysis between lung and kidney samples. Despite the large differences in the number of taxa identified, there was still a small number of overlapping species identified across the results of all software. We also found similarities in the software-overlapped DA taxa in lung vs kidney and in lung vs. spleen analyses, where two *Bordetella* species and a *Mycoplasm* species were reported by all software in both analyses. More DA identified were overlapped across software at the Phylum level. In addition to the overlapped DA taxa, Kaiju and Centrifuge, which are both index-based software, were more likely to report more numbers of taxa as differentially abundant than the other software. This two software were also the only two that reported both viral and archaeal taxa as DA (BLASTN only reported archaeal taxa and CLARK only reported virual taxa). Diamond was found to be the least sensitive in the DA analyses for all three comparisons between tissue samples, where phylum taxa identified by all the other software were frequently not identified by Diamond.

### Discrepancies in *Leptospira* Detection

For comparisons in sensitivity to identify the presence of the zoonotic pathogen *Leptospira* in all of our tissue samples, Centrifuge and Kaiju were found as the most sensitive software in diagnosing *Leptospira*, where Centrifuge reported the presence of *Leptospria* in all the samples. Since *Leptospira* colonizes the kidney of rats (Adler and de la Peña Moctezuma, 2015), we compared the results from three traditional methods (PCR/DFA/Culture) applied to kidney samples reported in a previous study (Rajeev *et al*., 2020). We found that most software included in our analysis had similar sensitivity in *Leptospira* identification with traditional methods, except for PCR. In addition, Centrifuge reported the presence of *Leptospira* in samples that were not reported by any other software or a traditional method. This identification could be due to Centrifuge’s better performance in sensitivity, or as a result of false positive reporting. Furthermore, we found that Kraken2 with maxikraken DB also reported *Leptospira*’s presence in all samples.

In conclusion, our study found that alignment-based software does not necessarily have better sensitivity in microbial profiling than alignment-free software. Diamond, one of the alignment-based software included in our analyses, reported the lowest sensitivity in DA analyses compared to other software. However, within the alignment-free software included in this study, two index-based software, Centrifuge and Kaiju, were found to be more sensitive than other software in microbial profiling, DA analysis, and pathogen detection. Metaphlan3, developed with a marker-based alignment-free algorithm, was found to have the lowest sensitivity in all the analyses when compared to all the other software included in this study. For the microbial community analyses, the characterization of within-samples microbial richness was largely impacted by software selection, but the impact was less significant if the characterization index used species abundance to weigh the index. A similar observation was found in the microbial community characterization analyses with different DBs. The within-sample richness characterization was mostly consistent when weighed by the species abundances within a sample. In addition, the presence of host genomes in the DBs does not largely impact the microbial profiling, but the overall compositions of microbial genomes included in the DBs impact the microbial classification the most. Decrease in the composition of microbial genomes in the DBs will largely decrease the sensitivity of the microbial classification. Moreover, we also found that the selection of the DBs can impact the ability of pathogen detection.

The inconsistencies found between the results of different metagenomic software showed that results obtained from metagenomic profiling analyses have the potential to be only the artifacts of the software’ algorithms. Shotgun metagenomics sequences might be too short for current taxonomical profiling software to differentiate microbial taxonomies between similar genomes (Tran and Phan, 2020). The use of wildly collected datasets have the advantage of addressing this challenge, reminding the investigators to stay skeptical with the classification results obtained from the profiling software. On the other hand, benchmarking the software’ performances with the *in vivo* dataset, in contrast to using *in silico* datasets, has the limitation of lacking knowledge about the true microbial compositions within each sample, which means we could not evaluate the performance of software based on their degrees of accuracy and sensitivity, nor giving direct suggestions on software’ selection. In addition, metagenomics profiling has been broadly used in many fields of studies, including clinical, pharmaceutical, as well as ecological. Each field uses microbial profiles differently based on the biological question proposed. Our choice of the wildly collected *Rattus* dataset could only address a limited number of software selection biases. We suggest researchers from different study fields to be aware of the possible error-prone conclusions made from metagenomics profiling analysis and evaluate it objectively comparing it to other traditional methods (e.g., PCR, culture, or serotyping). Advancement in sequencing as well as computational technologies allows modern-day biological research to move to a brand-new era. However, while benefiting from the powerfulness and convenience of technologies, we should always critically analyze and validate software outputs based on our prior knowledge and available evidence.

## Supporting information

Supplementary_FiguresS1-S4

Supplemental TableSI.kraken2_db_comparison

Supplemental TableSII.software_comparison

Supplemental TableSIII.lepto.diagnostic

Supplemental TableSIV.lepto.kraken2.db.diagnostic

## Acknowledgements

The sequence analysis work was supported by the National Science Foundation under Grant No. DGE-1545433 to R.X. and startup funds to L.C.M.S. from the University of Georgia Office of Research. The sample collection, sequencing and analysis was done during S.R.’s tenure at the Ross University School of Veterinary Medicine, Saint Kitts and it was supported by internal grants from the Center for One Health and Tropical Medicine. We also would like to thank Dr. Kanae Shiokawa for her help with collection and processing of rat specimens.

## Conflicts of interest

No conflict of interest declared.

## Repositories

The raw sequence files (FASTQ) were submitted to the NCBI Sequence Read Archive under the Bioproject accession number: PRJNA717669. The individual isolates can be accessed under the following Biosample accession numbers: SAMN18507082 - SAMN18507091. All scripts for this publication are freely available on the following Github link: https://github.com/rx32940/Metagenomics_tools

## Data summary

The raw sequence files (FASTQ) were submitted to the NCBI Sequence Read Archive under the Bioproject accession number: PRJNA717669. The individual isolates can be accessed under the following Biosample accession numbers: SAMN18507082 - SAMN18507091. The short-read archive accession numbers are listed in Table S1.

## Ethical Approval

Rats were captured following protocols approved by the Ross University School of Veterinary Medicine (RUSVM) IACUC (approval # 17-01-04).

## Supporting Information

SI.kraken2_db_comparison.xlsx

SII.software_comparison_full.xlsx

SIII.lepto.diagnostic.xlsx

SIV.lepto.kraken2.db.diagnostic.xlsx

JAM_Metagenomics_supplementary_figures.docx

Journal of Applied Microbiology_supplementary_text.docx

---

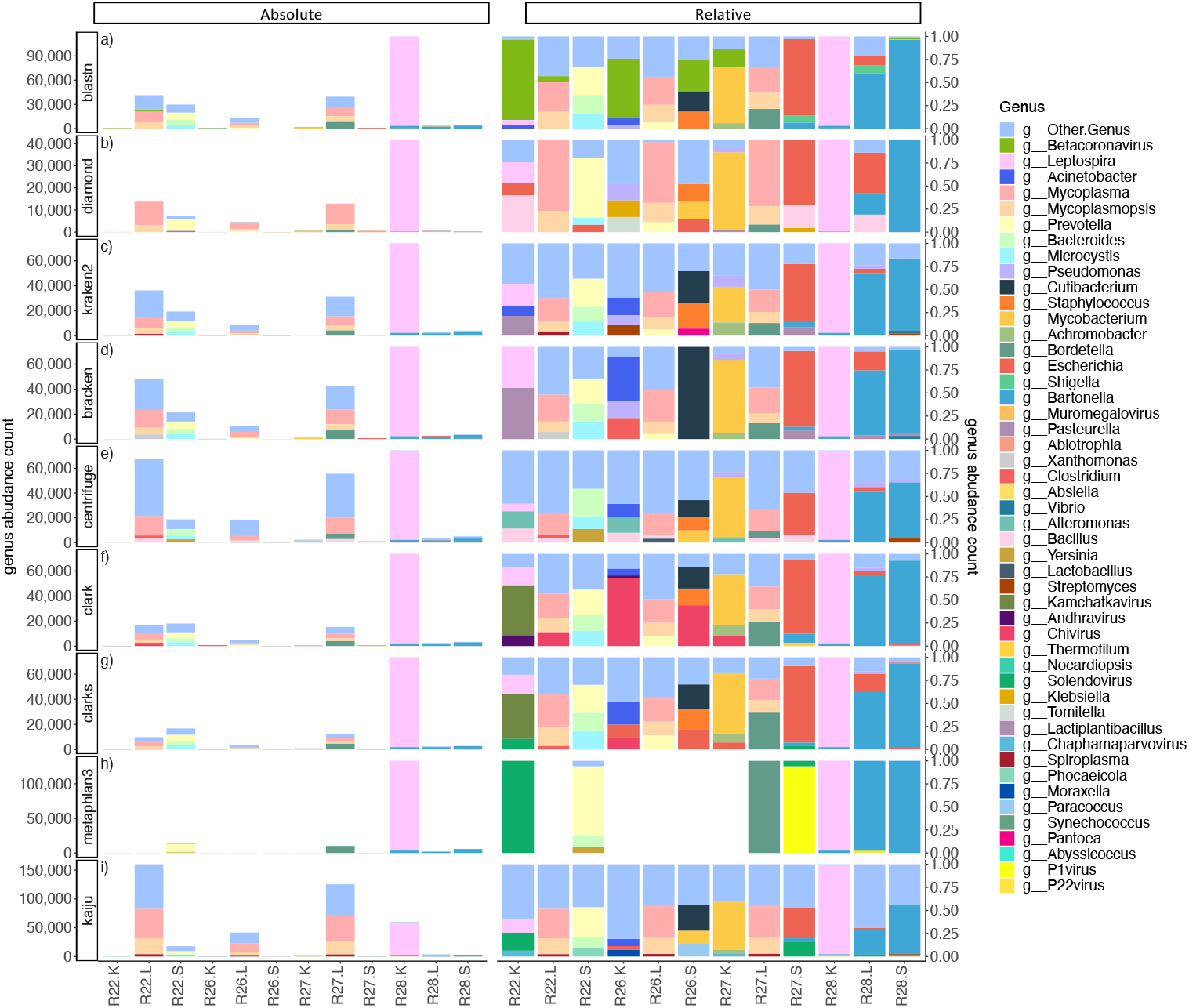

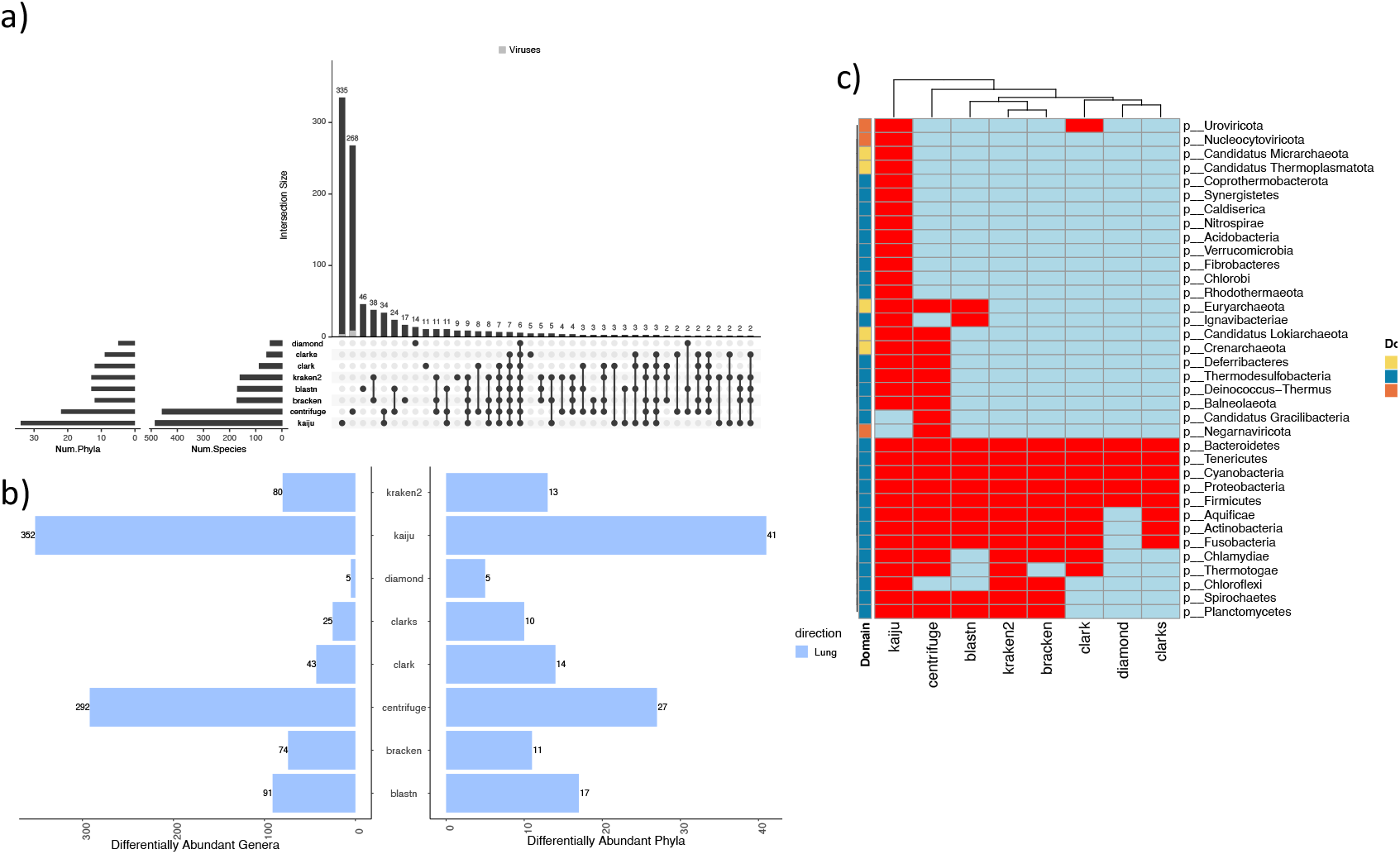

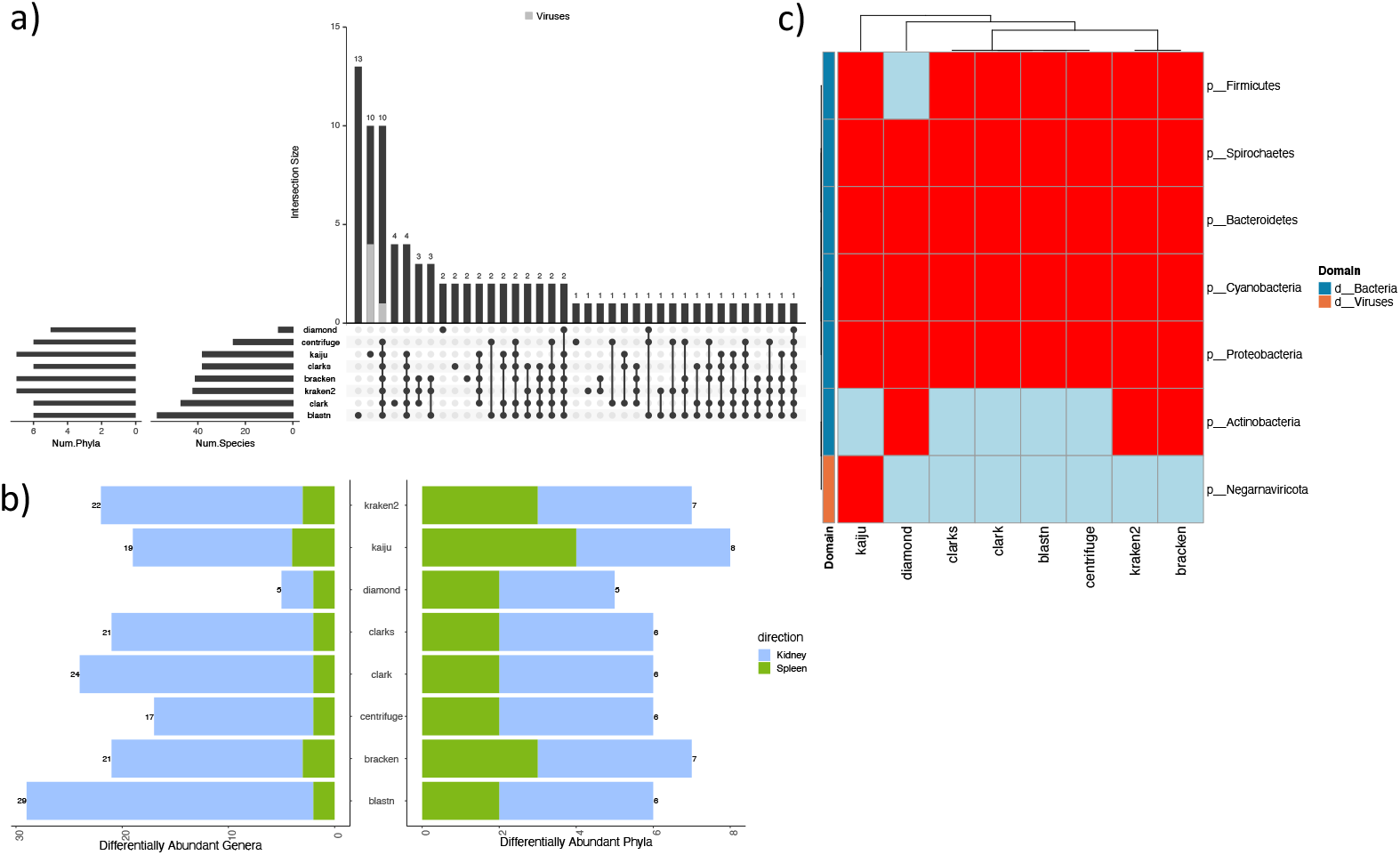

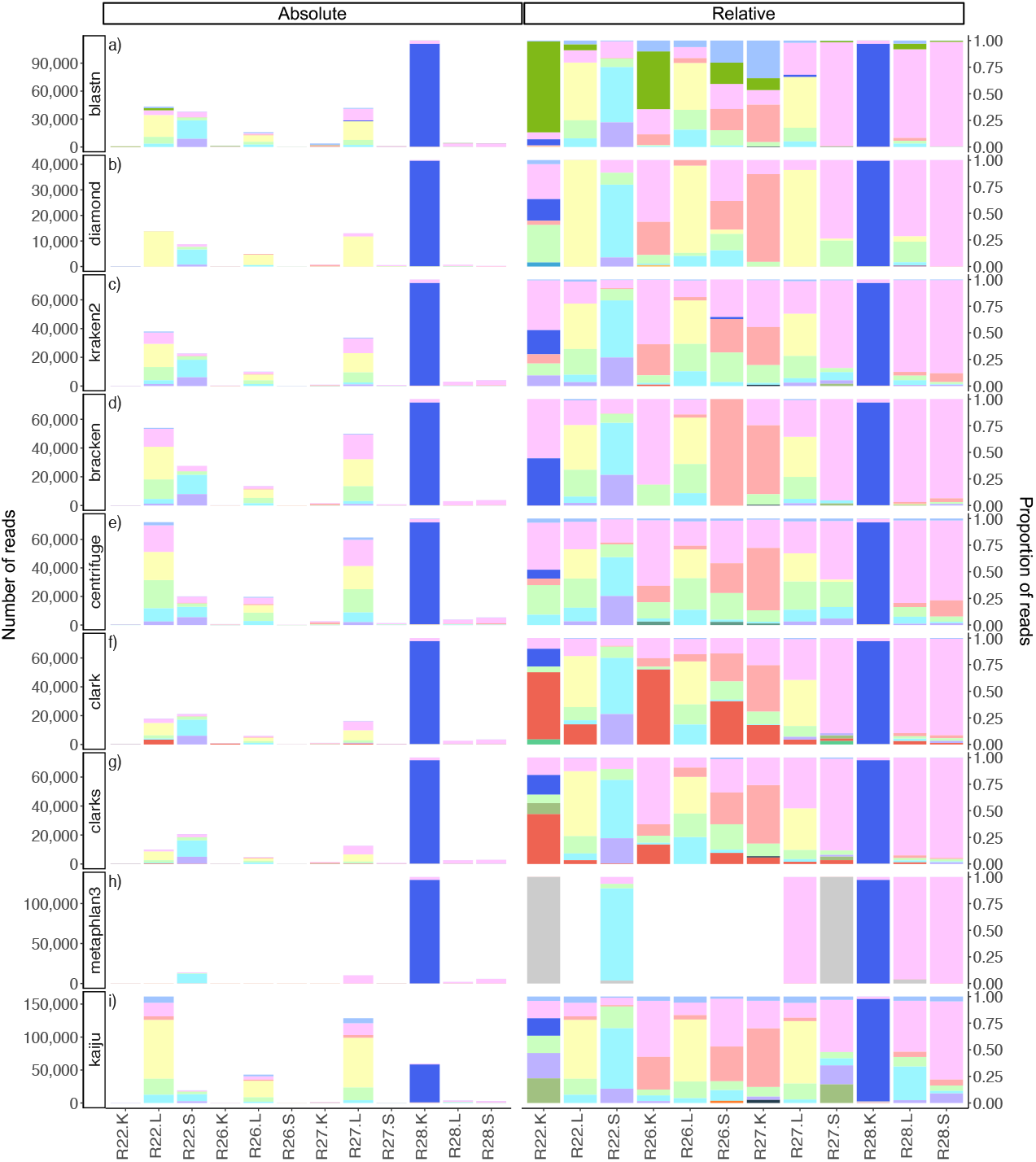

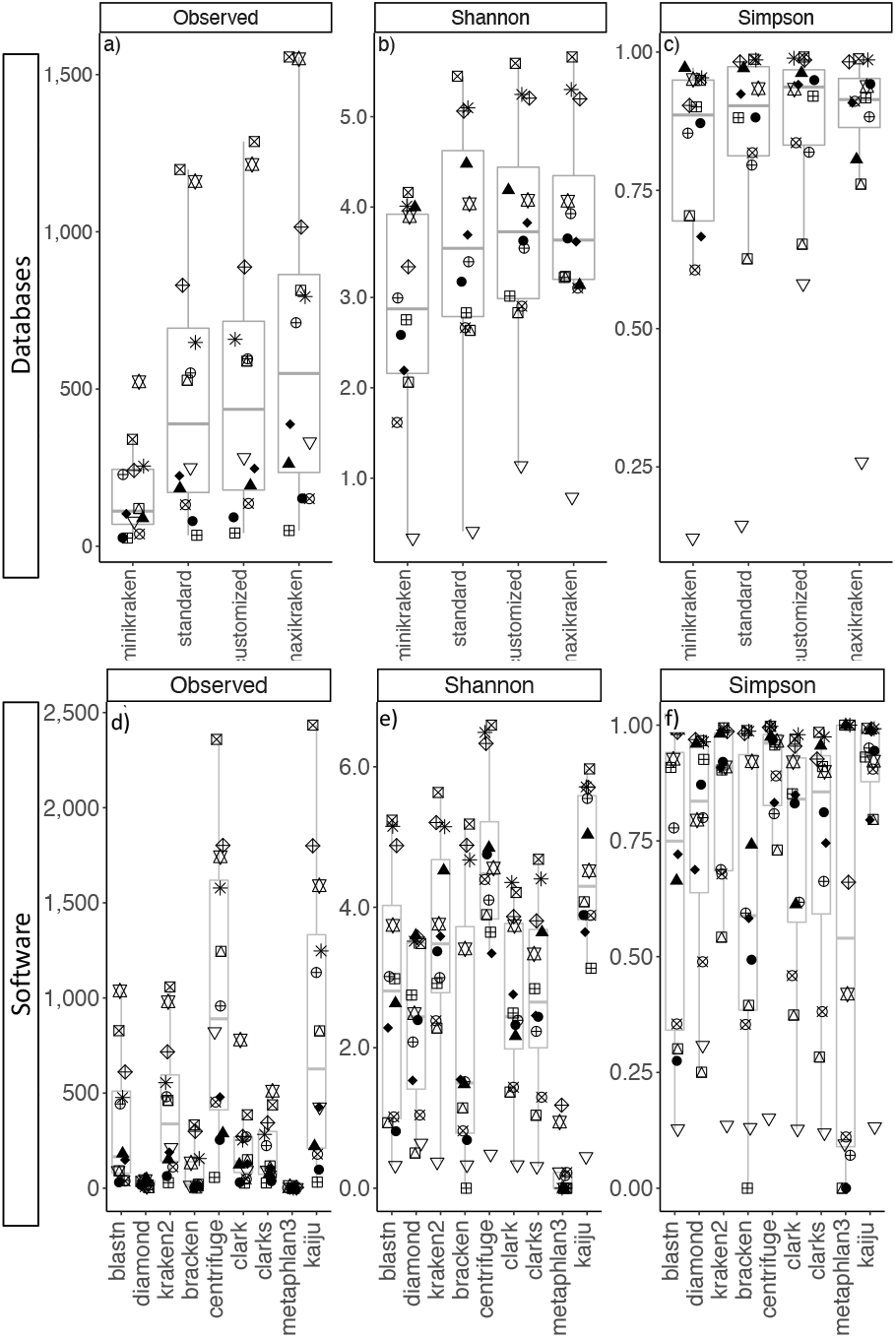

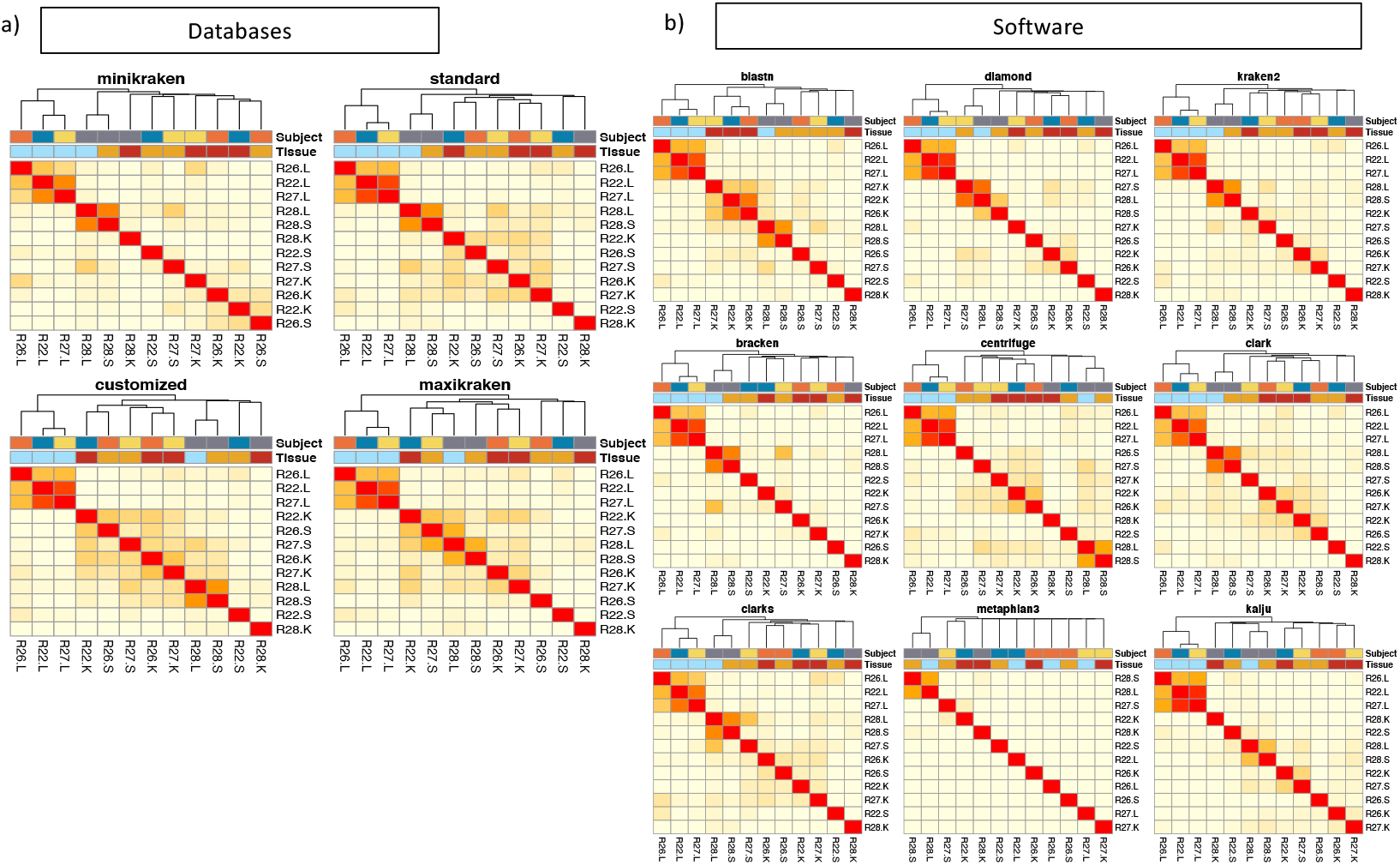

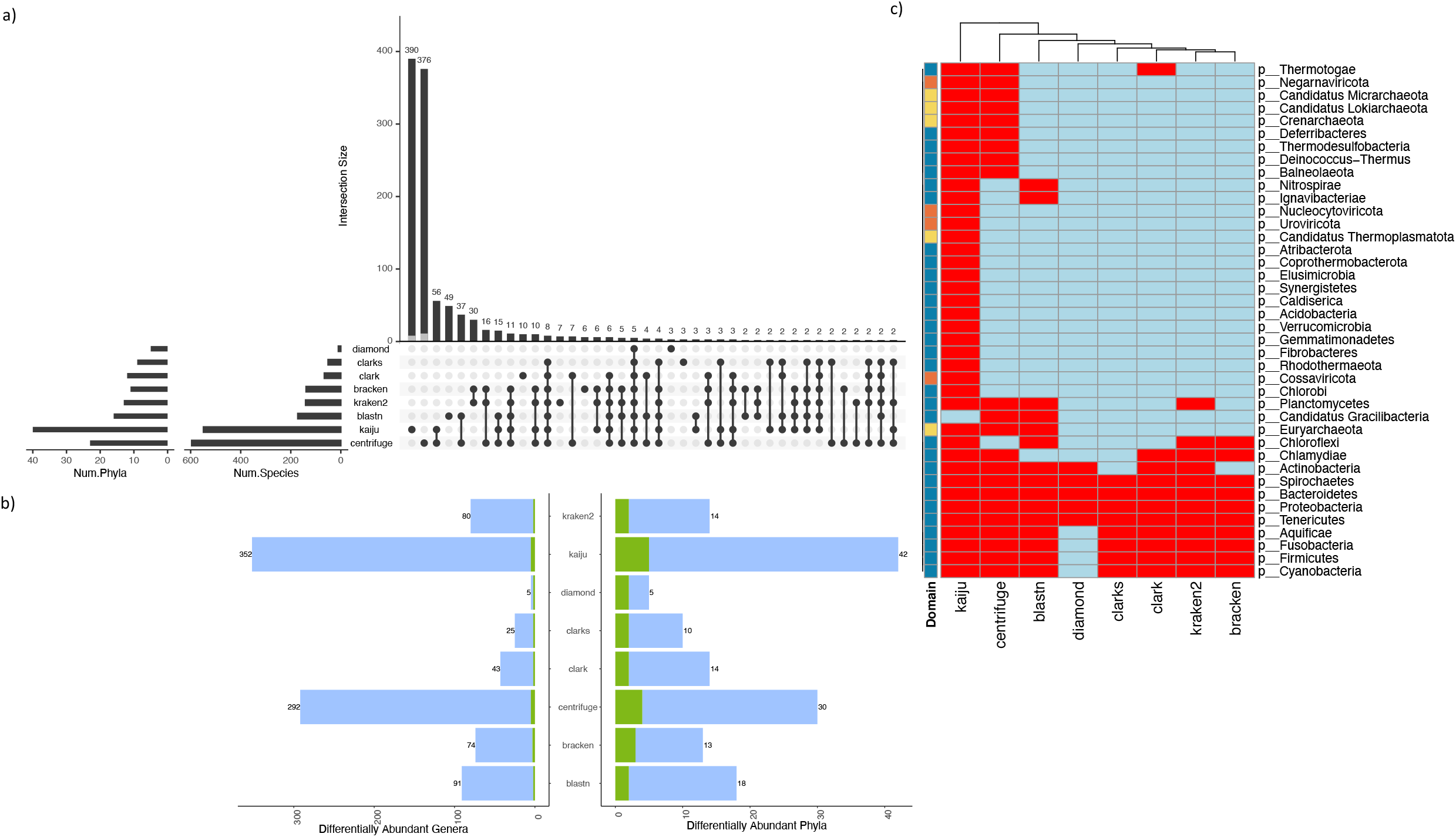

## Notes

https://github.com/rx32940/Metagenomics_tools

### Competing Interest Statement

The authors have declared no competing interest.

## References

Adler, B. and de la Peña Moctezuma (2015) ‘Leptospira and Leptospirosis’, Veterinary Microbiology, 140(3), pp. 287–296. doi:10.1007/978-3-662-45059-8.

Altschul, S.F. et al. (1990) ‘Basic local alignment search tool’, Journal of Molecular Biology, 215(3), pp. 403–410. doi:10.1016/S0022-2836(05)80360-2.

Ames, S.K. et al. (2015) ‘Using populations of human and microbial genomes for organism detection in metagenomes’, Genome Research, 25(7), pp. 1056–1067. doi:10.1101/gr.184879.114.

Beghini, F. et al. (2021) ‘Integrating taxonomic, functional, and strain-level profiling of diverse microbial communities with bioBakery 3’, eLife. Edited by P. Turnbaugh, E. Franco, and C.T. Brown, 10, p. e65088. doi:10.7554/eLife.65088.

Bolger, A.M., Lohse, M. and Usadel, B. (2014) ‘Trimmomatic: A flexible trimmer for Illumina sequence data’, Bioinformatics, 30(15). doi:10.1093/bioinformatics/btu170.

Bray, J.R. and Curtis, J.T. (1957) ‘An Ordination of the Upland Forest Communities of Southern Wisconsin’, Ecological Monographs, 27(4), pp. 325–349. doi:https://doi.org/10.2307/1942268.

Breitwieser, F.P., Lu, J. and Salzberg, S.L. (2019) ‘A review of methods and databases for metagenomic classification and assembly’, Briefings in Bioinformatics, 20(4), pp. 1125–1136. doi:10.1093/bib/bbx120.

Buchfink, B., Xie, C. and Huson, D.H. (2015) ‘Fast and sensitive protein alignment using DIAMOND’, Nature Methods, 12(1), pp. 59–60. doi:10.1038/nmeth.3176.

Burrows, M. and Wheeler, D.J. (1994) A block-sorting lossless data compression algorithm.

Camacho, C. et al. (2009) ‘BLAST+: Architecture and applications’, BMC Bioinformatics [Preprint]. doi:10.1186/1471-2105-10-421.

Cannings, C. (2004) ‘Mathematical and Statistical Methods for Genetic Analysis (2nd ed)’, Heredity, 92(1), pp. 51–51. doi:10.1038/sj.hdy.6800368.

Chavira, A. et al. (2019) ‘The Microbiome and Its Potential for Pharmacology’, Concepts and Principles of Pharmacology: 100 Years of the Handbook of Experimental Pharmacology. Edited by J.E. Barrett, C.P. Page, and M.C. Michel, pp. 301–326. doi:10.1007/164_2019_317.

Chen, Y.-Y. et al. (2019) ‘Microbiome–metabolome reveals the contribution of gut–kidney axis on kidney disease’, Journal of Translational Medicine, 17(1), p. 5. doi:10.1186/s12967-018-1756-4.

Clark, D.P. and Pazdernik, N.J. (2016) ‘Environmental Biotechnology’, in Biotechnology. Elsevier, pp. 393–418. doi:10.1016/B978-0-12-385015-7.00012-0.

Desmonts, G. and Remington, J.S. (1980) ‘Direct agglutination test for diagnosis of Toxoplasma infection: method for increasing sensitivity and specificity’, Journal of Clinical Microbiology, 11(6), pp. 562–568. doi:10.1128/jcm.11.6.562-568.1980.

Driscoll, J.R. (2009) ‘Spoligotyping for molecular epidemiology of the Mycobacterium tuberculosis complex’, Methods in Molecular Biology (Clifton, N.J.), 551, pp. 117–128. doi:10.1007/978-1-60327-999-4_10.

Durazzi, F. et al. (2021) ‘Comparison between 16S rRNA and shotgun sequencing data for the taxonomic characterization of the gut microbiota’, Scientific Reports, 11(1), p. 3030. doi:10.1038/s41598-021-82726-y.

Escobar-Zepeda, A. et al. (2018) ‘Analysis of sequencing strategies and tools for taxonomic annotation: Defining standards for progressive metagenomics’, Scientific Reports, 8(1), p. 12034. doi:10.1038/s41598-018-30515-5.

Fouhy, F. et al. (2016) ‘16S rRNA gene sequencing of mock microbial populations-impact of DNA extraction method, primer choice and sequencing platform’, BMC Microbiology, 16(1), p. 123. doi:10.1186/s12866-016-0738-z.

Galbraith, D.A. et al. (2018) ‘Investigating the viral ecology of global bee communities with high-throughput metagenomics’, Scientific Reports, 8(1), p. 8879. doi:10.1038/s41598-018-27164-z.

Ghosh, A., Mehta, A. and Khan, A.M. (2019) ‘Metagenomic Analysis and its Applications’, in Ranganathan, S. et al. (eds). Oxford: Academic Press, pp. 184–193. doi:https://doi.org/10.1016/B978-0-12-809633-8.20178-7.

Ginestet, C. (2011) ‘ggplot2: Elegant Graphics for Data Analysis’, Journal of the Royal Statistical Society: Series A (Statistics in Society) [Preprint]. doi:10.1111/j.1467-985x.2010.00676_9.x.

Granjou, C. and Phillips, C. (2019) ‘Living and labouring soils: Metagenomic ecology and a new agricultural revolution?’, BioSocieties, 14(3). doi:10.1057/s41292-018-0133-0.

Grossart, H.-P. et al. (2020) ‘Linking metagenomics to aquatic microbial ecology and biogeochemical cycles’, Limnology and Oceanography, 65(S1). doi:10.1002/lno.11382.

Grützke, J. et al. (2021) ‘Direct identification and molecular characterization of zoonotic hazards in raw milk by metagenomics using Brucella as a model pathogen’, Microbial Genomics, 7(5), p. 000552. doi:10.1099/mgen.0.000552.

Handelsman, J. (2004) ‘Metagenomics: Application of Genomics to Uncultured Microorganisms’, Microbiology and Molecular Biology Reviews, 68(4), pp. 669–685. doi:10.1128/MMBR.68.4.669-685.2004.

Handley, K.M. (2019) ‘Determining Microbial Roles in Ecosystem Function: Redefining Microbial Food Webs and Transcending Kingdom Barriers’, mSystems, 4(3). doi:10.1128/mSystems.00153-19.

Healy, J. and Chambers, D. (2014) ‘Approximate $k$-Mer Matching Using Fuzzy Hash Maps’, IEEE/ACM Transactions on Computational Biology and Bioinformatics, 11(1), pp. 258–264. doi:10.1109/TCBB.2014.2309609.

Holm, S. (1979) ‘A Simple Sequentially Rejective Multiple Test Procedure’, Scandinavian Journal of Statistics, 6(2), pp. 65–70. Available at: https://www.jstor.org/stable/4615733 (Accessed: 27 February 2022).

Janda, J.M. and Abbott, S.L. (2007) ‘16S rRNA gene sequencing for bacterial identification in the diagnostic laboratory: Pluses, perils, and pitfalls’, Journal of Clinical Microbiology. American Society for Microbiology Journals, pp. 2761–2764. doi:10.1128/JCM.01228-07.

Johnson, J.S. et al. (2019) ‘Evaluation of 16S rRNA gene sequencing for species and strain-level microbiome analysis’, Nature Communications, 10(1), p. 5029. doi:10.1038/s41467-019-13036-1.

Johnson, M. et al. (2008) ‘NCBI BLAST: a better web interface.’, Nucleic acids research [Preprint]. doi:10.1093/nar/gkn201.

Jovel, J. et al. (2016) ‘Characterization of the Gut Microbiome Using 16S or Shotgun Metagenomics’, Frontiers in Microbiology, 7. doi:10.3389/fmicb.2016.00459.

Kassambara, A. (2021) rstatix: Pipe-Friendly Framework for Basic Statistical Tests. [R package version 0.7.0]. Available at: https://CRAN.R-project.org/package=rstatix.

Kim, D. et al. (2016) ‘Centrifuge: rapid and sensitive classification of metagenomic sequences’, Genome Research, 26(12), pp. 1721–1729. doi:10.1101/gr.210641.116.

Knights, D., Lassen, K.G. and Xavier, R.J. (2013) ‘Advances in inflammatory bowel disease pathogenesis: linking host genetics and the microbiome’, Gut, 62(10), pp. 1505–1510. doi:10.1136/gutjnl-2012-303954.

Langmead, B. et al. (2019) ‘Scaling read aligners to hundreds of threads on general-purpose processors’, Bioinformatics, 35(3), pp. 421–432. doi:10.1093/bioinformatics/bty648.

Lequin, R.M. (2005) ‘Enzyme Immunoassay (EIA)/Enzyme-Linked Immunosorbent Assay (ELISA)’, Clinical Chemistry, 51(12), pp. 2415–2418. doi:10.1373/clinchem.2005.051532.

Love, M.I., Huber, W. and Anders, S. (2014) ‘Moderated estimation of fold change and dispersion for RNA-seq data with DESeq2’, Genome Biology, 15(12), p. 550. doi:10.1186/s13059-014-0550-8.

Lu, J.et al. (2017) ‘Bracken: estimating species abundance in metagenomics data’, PeerJ Computer Science, 3, p. e104. doi:10.7717/peerj-cs.104.

Mashiane, R.A. et al. (2017) ‘Metagenomic analyses of bacterial endophytes associated with the phyllosphere of a Bt maize cultivar and its isogenic parental line from South Africa’, World Journal of Microbiology and Biotechnology, 33(4). doi:10.1007/s11274-017-2249-y.

McMurdie, P.J. and Holmes, S. (2013) ‘phyloseq: An R Package for Reproducible Interactive Analysis and Graphics of Microbiome Census Data’, PLOS ONE, 8(4), p. e61217. doi:10.1371/journal.pone.0061217.

Menzel, P., Ng, K.L. and Krogh, A. (2016) ‘Fast and sensitive taxonomic classification for metagenomics with Kaiju’, Nature Communications, 7(1), p. 11257. doi:10.1038/ncomms11257.

Oksanen, J. et al. (2013) ‘Package vegan’, R Packag ver [Preprint].

Ounit, R. et al. (2015) ‘CLARK: fast and accurate classification of metagenomic and genomic sequences using discriminative k-mers’, BMC Genomics [Preprint]. doi:10.1186/s12864-015-1419-2.

Ounit, R. and Lonardi, S. (2016) ‘Higher classification sensitivity of short metagenomic reads with CLARK-S’, Bioinformatics [Preprint]. doi:10.1093/bioinformatics/btw542.

Peabody, M.A. et al. (2015) ‘Evaluation of shotgun metagenomics sequence classification methods using in silico and in vitro simulated communities’, BMC Bioinformatics, 16(1), p. 362. doi:10.1186/s12859-015-0788-5.

Pereira-Marques, J. et al. (2019a) ‘Impact of Host DNA and Sequencing Depth on the Taxonomic Resolution of Whole Metagenome Sequencing for Microbiome Analysis’, Frontiers in Microbiology, 10. doi:10.3389/fmicb.2019.01277.

Pereira-Marques, J. et al. (2019b) ‘Impact of Host DNA and Sequencing Depth on the Taxonomic Resolution of Whole Metagenome Sequencing for Microbiome Analysis’, Frontiers in Microbiology, 10. doi:10.3389/fmicb.2019.01277.

Qin, J. et al. (2012) ‘A metagenome-wide association study of gut microbiota in type 2 diabetes’, Nature, 490(7418), pp. 55–60. doi:10.1038/nature11450.

R Core Team (2020) ‘R: A Language and Environment for Statistical Computing’, R Foundation for Statistical Computing [Preprint]. Available at: https://www.r-project.org/ (Accessed: 25 March 2021).

Rajeev, S. et al. (2020) ‘Detection and Characterization of Leptospira Infection and Exposure in Rats on the Caribbean Island of Saint Kitts’, Animals, 10(2), p. 350. doi:10.3390/ani10020350.

Ranjan, R. et al. (2016) ‘Analysis of the microbiome: Advantages of whole genome shotgun versus 16S amplicon sequencing’, Biochemical and Biophysical Research Communications, 469(4), pp. 967–977. doi:10.1016/J.BBRC.2015.12.083.

Shannon, C.E. (1948) ‘A Mathematical Theory of Communication’, m The Bell System Technical Journal, 27, pp. 379–423. doi:10.1002/j.1538-7305.1948.tb01338.x.

Sharpton, T.J. (2014) ‘An introduction to the analysis of shotgun metagenomic data’, Frontiers in Plant Science, 5. doi:10.3389/fpls.2014.00209.

Simpson, E.H. (1949) ‘Measurement of Diversity’, Nature, 163(4148), pp. 688–688. doi:10.1038/163688a0.

Skarżyńska, M. et al. (2020) ‘A metagenomic glimpse into the gut of wild and domestic animals: Quantification of antimicrobial resistance and more’, PLOS ONE, 15(12), p. e0242987. doi:10.1371/journal.pone.0242987.

The Huttenhower Lab (no date) KneadData. Available at: https://huttenhower.sph.harvard.edu/kneaddata/ (Accessed: 25 March 2021).

Tran, Q. and Phan, V. (2020) ‘Assembling Reads Improves Taxonomic Classification of Species’, Genes, 11(8), p. 946. doi:10.3390/genes11080946.

Truong, D.T. et al. (2015) ‘MetaPhlAn2 for enhanced metagenomic taxonomic profiling’, Nature Methods, 12(10), pp. 902–903. doi:10.1038/nmeth.3589.

Tun, H.M. et al. (2012) ‘Gene-centric metagenomics analysis of feline intestinal microbiome using 454 junior pyrosequencing’, Journal of Microbiological Methods, 88(3), pp. 369–376. doi:10.1016/j.mimet.2012.01.001.

Wang, J.-J. et al. (2019) ‘Metagenomic analysis of gut microbiota alteration in a mouse model exposed to mycotoxin deoxynivalenol’, Toxicology and Applied Pharmacology, 372, pp. 47–56. doi:10.1016/j.taap.2019.04.009.

Whittaker, R.H. (1960) ‘Vegetation of the Siskiyou Mountains, Oregon and California’, Ecological Monographs, 30(3), pp. 279–338. doi:https://doi.org/10.2307/1943563.

Woese, C.R., Kandlert, O. and Wheelis, M.L. (1990) ‘Towards a natural system of organisms: Proposal for the domains Archaea, Bacteria, and Eucarya’, Proc. Nati. Acad. Sci. USA, 87, pp. 4576–4579. doi:10.1073/pnas.87.12.4576.

Wood, D.E., Lu, J. and Langmead, B. (2019) ‘Improved metagenomic analysis with Kraken 2’, Genome Biology [Preprint]. doi:10.1186/s13059-019-1891-0.

Yang, S. and Rothman, R.E. (2004) ‘PCR-based diagnostics for infectious diseases: uses, limitations, and future applications in acute-care settings’, The Lancet. Infectious Diseases, 4(6), pp. 337–348. doi:10.1016/S1473-3099(04)01044-8.

Ye, S.H. et al. (2019) ‘Benchmarking Metagenomics Tools for Taxonomic Classification.’, Cell, 178(4), pp. 779–794. doi:10.1016/j.cell.2019.07.010.

Zhong, H. et al. (2019) ‘Distinct gut metagenomics and metaproteomics signatures in prediabetics and treatment-naïve type 2 diabetics’, EBioMedicine, 47, pp. 373–383. doi:10.1016/j.ebiom.2019.08.048.

Zielezinski, A. et al. (2017) ‘Alignment-free sequence comparison: benefits, applications, and tools’, Genome Biology, 18(1), p. 186. doi:10.1186/s13059-017-1319-7.

