## Supplementary_TextS1-S3 for "Selection of software and database for metagenomics sequence analysis impacts the outcome of microbial profiling and pathogen detection"

**Supplementary Text 1. Resources for Database (DB) setup**

The DBs version numbers and compositions used for each software, as well as the computational resources and time used to build them, are described in Table 1. For Kraken 2, four DBs were used: maxikraken 2, standard, minikraken, and customized. The maxikraken2 DB includes both complete and draft genomes of the bacteria, archaea, fungi, protozoa and viruses, thus requires over 150 GB of memory available on the workstation for the downstream analyses, while the standard and minikraken DBs require only 53 GB and 8GB, respectively. The customized DB took 60 GB and it was built with the same composition of the standard DB and with the addition of the genomes of the two *Rattus* species used in this study. All analyses were run in the high-performance computing cluster at the Georgia Advanced Computing Resource Center. With 12 threads of CPU used on the high memory computing node, the building of the customized DB took ~15 hrs (Table I). The time of each analysis changed with the selection of the DB used, but the time differences were small (only in the range of minutes). For the rest of the software, pre-built DBs were chosen to perform the profiling, if these were provided by the software. CLARK, CLARK-s, Diamond, and Kaiju were the only four software in our study without a pre-built software provided. With 12 threads of CPU used on the high memory computing node, the building of CLARK’s DB took over 42 hours to complete, with over 400 GB memory used (Table I). CLARK-s DB, required to build on top of the CLARK’s DB, took around 40 additional hours to complete, with around 300 GB memory used (Table I). The building of Diamond’s DB, with the same computational setting, completed in ~2.4 hours using ~ 8 GB, while Kaiju’s DB took ~ 5 hours to complete using ~115 GB of memory (Table I). As for analysis time, using 12 threads of CPU on the UGA’s high memory computing node, Diamond used on average ~5 hours to classify one sample, while BLASTN used ~2 hours. The rest of the software classified one sample within 5 minutes at most (Table I).

**Supplementary Text2. DB comparisons at genus and species levels.**

For the Eukaryota taxon, all but the number of reads classified by the standard and customized DBs were found not to have statistically significant differences (Figure 1a). However, the classification profiles of these DBs at the genus level assigned different number of Eukaryota reads to the two hosts (“g__Rattus” and “g__Homo”). The standard DB profiling assigned all 9,771 Eukaryota reads to “p__Homo” while the customized DB profiling assigned only 4,941 Eukaryota reads to “p__Homo” and 4,961 to “g__Rattus” (Table SI.1, Figure S1b-c). Minikraken DB assigned only 986 to the Eukaryota taxon and all its reads were assigned to “g__Homo” (Table SI.1, Figure S1a), while maxikraken DB assigned 5,527 reads under the Eukaryota taxon, with 153 unique genus from human, fungi (ex. *Pneumocystis murina*), ciliate (*Paramecium tetraurelia*), amoebae (*Naegleria gruberi*), parasites (ex. *Plasmodium vivax*), etc. For the Bacteria taxon, all the pairwise comparisons of the number of reads classified by the different DBs were found to have statistically significant differences (Figure 1b), which could be further validated by Bacteria taxon’s classification at the genus and species levels. For example, the genus taxa “g__Enterococcus” (species: *E. faecium*) and “g__Bacillus” (species: *B. sp. SRB_28 & B. sp. SRB_336* ) were identified as the top two most abundant bacteria taxa in maxikraken DB’s classification profile for sample R22.K (Figure S1d), but “g__Enterococcus” taxon was not identified by any other DBs’ classification, and only 5 reads were identified by the minikraken DB as “g__Bacillus” (species: *B. megaterium*) (Figure S1a-c). Similar observations with high abundance of “g__Enterococcus” and “g__Bacillus genus taxa were identified by maxikraken DB’s classification but not by any other DBs that were also found in the classification profiles of R26.K, R26.S, R27.K, R27.S, and R28.L (Figure S1). In addition, the genus taxa “g__Prevotella” were identified as the most abundant taxa in the sample R22.S by standard (5,954), customized (6,288), and maxikraken (8,815) DBs’ classification, mostly from species *P. copri* (5,869; 6,142; 6,511 reads, respectively) (Figure S1b-d), while minikraken identified in total 218 reads from 11 species of genus “g__Prevotella” taxon, but none from species *P. copri* (Figure S1a). For reads classified under the Viruses taxon, only the comparisons standard vs customized, and minikraken vs maxikraken DBs were found not to have statistically significant differences. When looking at Viruses classifications at the genus level, 10 (SD 17), 33 (SD 50), 37 (SD 54), and 10 (SD 22) genus (species) level taxa were identified by minikraken, standard, and maxikraken DBs, respectively (Table SI). Genus taxa “g__Alphabaculovirus” (species: Adoxophyes orana nucleopolyhedrovirus) and “g__Muromegalovirus” (species: Murid betaherpesvirus 2) were identified as two of the most abundant Viruses genus taxa by minikraken, standard, and customized DBs, but only Muromegalovirus was identified using maxikraken (Table SI). In addition, 91 and 112 reads of genus taxa “g__Andhravirus” (species: Staphylococcus virus Andhra) and 72 and 72 reads of “g__Alphanudivirus” (species: Oryctes rhinoceros nudivirus) were identified by standard and customized DBs as one of the top most abundant Viruses taxa, but these two taxa were not identified by minikraken nor maxikraken DBs’ (Table SI). In the case of Archaea, only the classification results of minikraken DB were found to have statistically significant differences when compared with the results of other DBs, and the classification results of the other three DBs did not have statistically significant differences between each other. However, when looking at the lower-level classifications for Archaea, 106, 247, 255, and 1,546 Archaea reads were classified under 4 (SD: 5), 34 (SD: 39), 38 (SD: 45), and 25 (SD: 28) unique genus (species) taxa by minikraken, standard, customized, and maxikraken DBs, respectively. The genus taxon “g__Methanobrevibacter” was identified as one of the most abundant ones across samples by all four DBs, however, minikraken, standard, and customized DB identified the species taxa *M.milerae* and *M. smithii* from the genus taxon, while maxikraken identified the species *M. oralis* and *M. filiformis* (Table SI). In addition, 1, 9, and 11 reads were classified under the genus taxon, “g__Methanococcus” (species: *M. maripaludis*) by minikraken, standard, and customized DBs, but 520 reads were classified under this taxon when using the maxikraken DB (Table SI).

**Supplementary Text3. Differentially Abundant (DA) analysis between lung and spleen samples and between kidney and lung samples.**

*Lung and spleen sample comparison*

The DA taxa identified between lung and spleen samples were similar with those identified between lung and kidney samples (Table SII.8, Figure S2). Kaiju identified the highest number of DA species (484 taxa), while Diamond identified the lowest (44 taxa) (Figure S2a). All the DA taxa were more abundant in the lung than in the spleen samples (Figure S2b). Six species (*Mycoplasm pulmonis*, *Mycoplasma bovoculi*, *Mycoplasma neurolyticum*, *Bordetella pseudohinzii*, *Bordetella bronchiseptica*, and *Bacteroides uniformis*) were identified as the DA taxa by all software (Table SII.8). Kaiju has the highest number of distinct DA species taxa (335), followed by centrifuge (268), and BLASTN (46) (Figure S2a).

At the Phylum level, “p__Bacterodietes”, “p__Tenericutes”, “p__Cyanobacteria”, “p__Protebacteria”, and “p__Firmicutes” were as DA identified by all the software (Figure S2c). Taxa "p__Aquificae”, "p__Actinobacteria”,and “p__Fusobacteria” were identified by all software except for Diamond. Archaea phylum, "p__Euryarchaeota”, was still the Archaea taxon identified by BLASTN, Centrifuge, and Kaiju, however, the rest of the Archaea taxa were either only identified by Kaiju and Centrifuge, or Kaiju alone. Virus taxon, “p__Negarnaviricota”, was only identified by Centrifuge as differentially abundant, while Kaiju only identified the virus taxa “p__Nucleocytoviricota” and “p__Uroviricota”. Morever, CLARK also reported the virus taxon, “p__Uroviricota”, as significantly abundant.

*Kidney and Spleen sample comparison*

The DA taxa between kidney and spleen samples are presented in Table SII.9 and Figure S3. The number of species identified ranges from 6 by Diamond and 57 by BLASTN (Figure S3a). More taxa were identified significantly abundant in the kidney samples than in the spleen ones, especially at the genus level (Figure S3b). Kaiju, the software that identified the second highest number of distinct DA taxa at the species level, has five out of ten distinct taxa reported as viruses (Figure S3a). In general, only one species (*Leptospira interrogans*) and four phylum taxa (“p__Spirochaetes”, “p__Bacteroidetes", "p__Cyanobacteria”, and “p__Proteobacteria”) were reported by all software (Table SII.9, Figure S3c). The Phylum taxon “p__Firmicutes” was identified as the DA taxon by all software except for Diamond. Kaiju solely identified the virus taxon, “p__Negarnaviricota”, as the DA taxon in this analysis.
