## Supplementary_FiguresS1-S4 for "Selection of software and database for metagenomics sequence analysis impacts the outcome of microbial profiling and pathogen detection"

**Supplementary Figures**

**Figure S1.** Genus-level microbial profiles for rat tissue samples using four different databases (minikraken, standard, customize, maxikraken) (a-d). Each row panel represent microbial profiles classified using a different database. Left panel is the microbial profiles reported in absolute number of reads and right panel is the microbial profiles reported in relative number of reads. Top 5 most abundant genus taxa for every sample are shown explicitly, while rest of the taxa were combined under “g__Other.Genus” in the figure below.
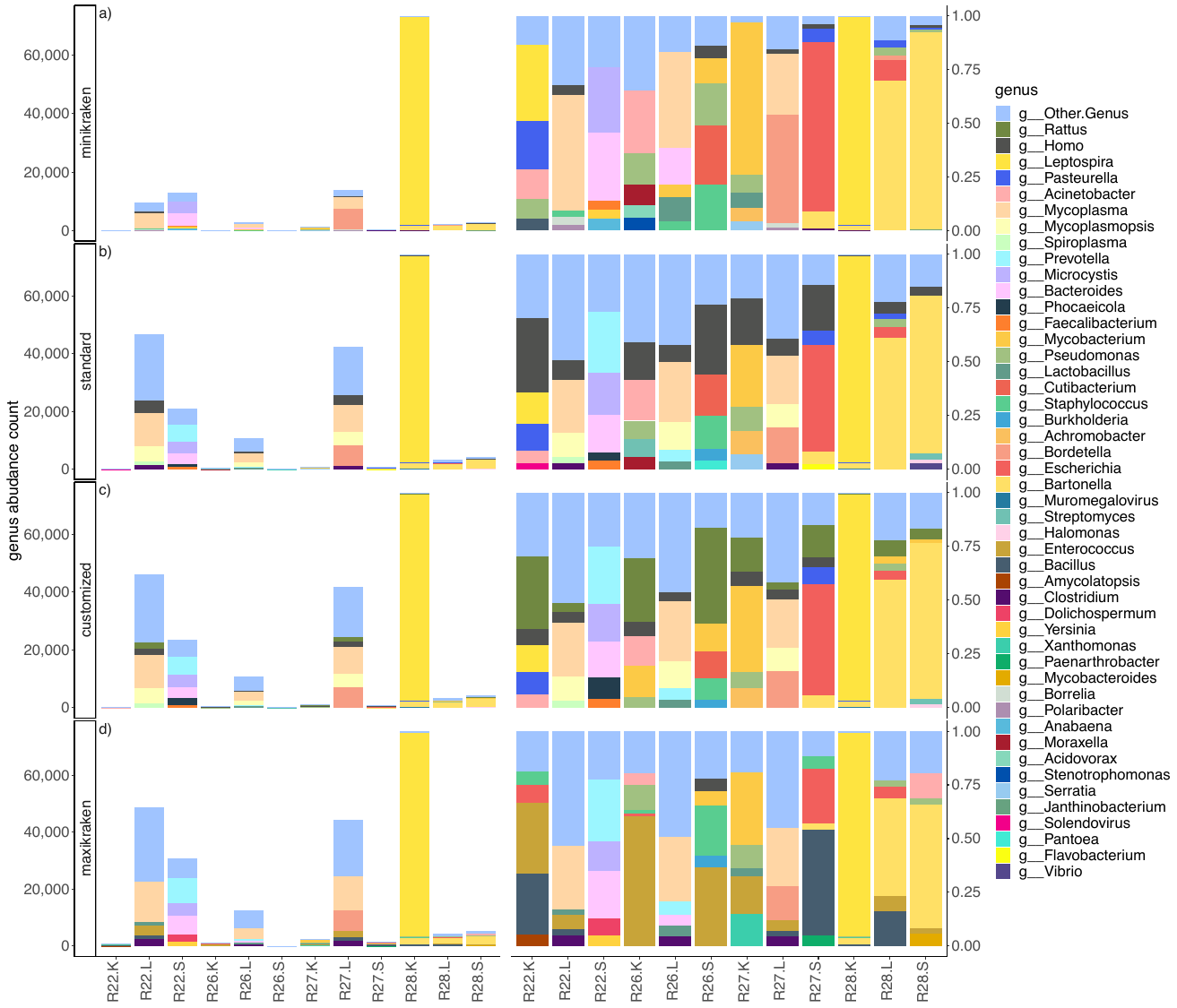

**Figure S2.** Genus-level microbial profiles for rat tissue samples using nine different software (a-i). Each row panel represent microbial profiles classified using a different software. Left panel is the microbial profiles reported in absolute number of reads and right panel is the microbial profiles reported in relative number of reads.

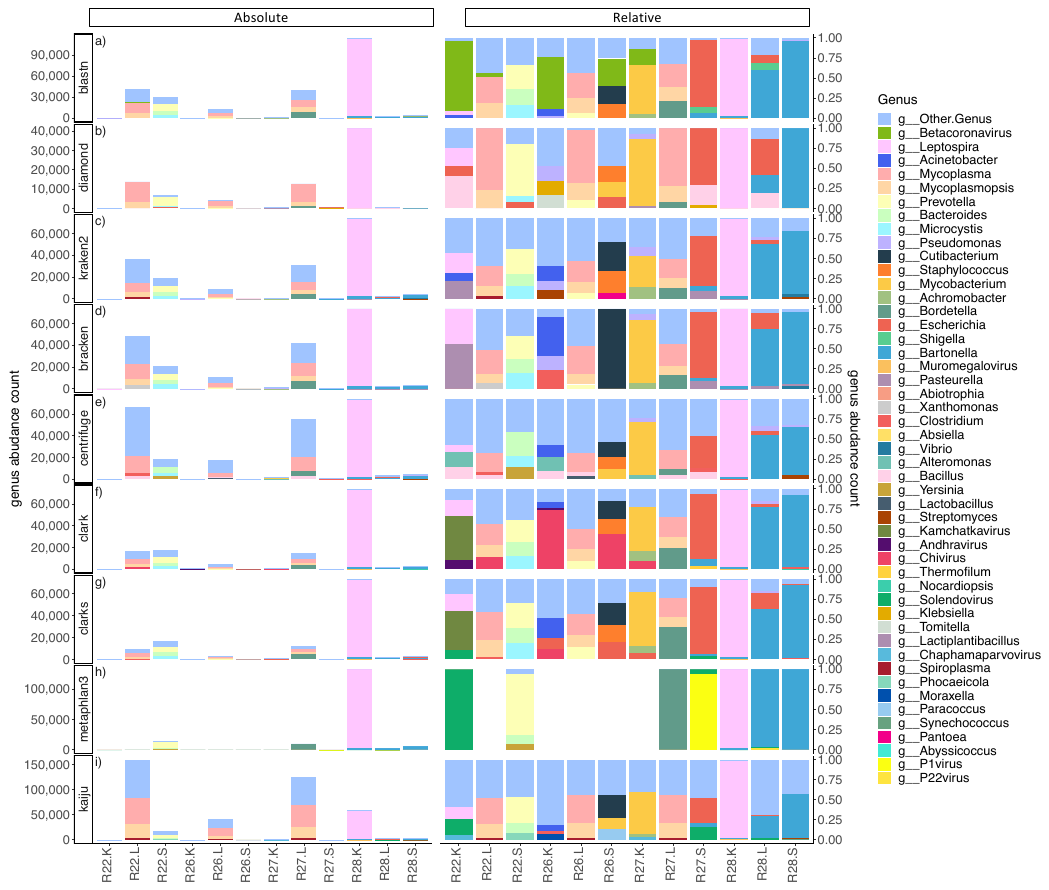

**Figure S3.** DA microbial taxa identified from comparing the microbial profiles of all rats’ lung and spleen tissues. The number of DA species and phylum identified using different software’s profile is shown at the bar plot directly left to the software names in a). The intersection between DA taxa identified by different software at the species level is shown at the barplot at the top of a), where the number of DA virus taxa were colored in gray. Dotplot at the bottom shown the combinations of intersections between software. Number of DA taxa at phylum and genus level is shown in b). The DA phylum taxa identified by each software is shown in c), where red (
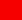
)indicate the phylum taxa at each row is reported as differentially abundant by the classification of the software in every column, and blue (
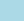
) is not reported as DA taxa by the software.

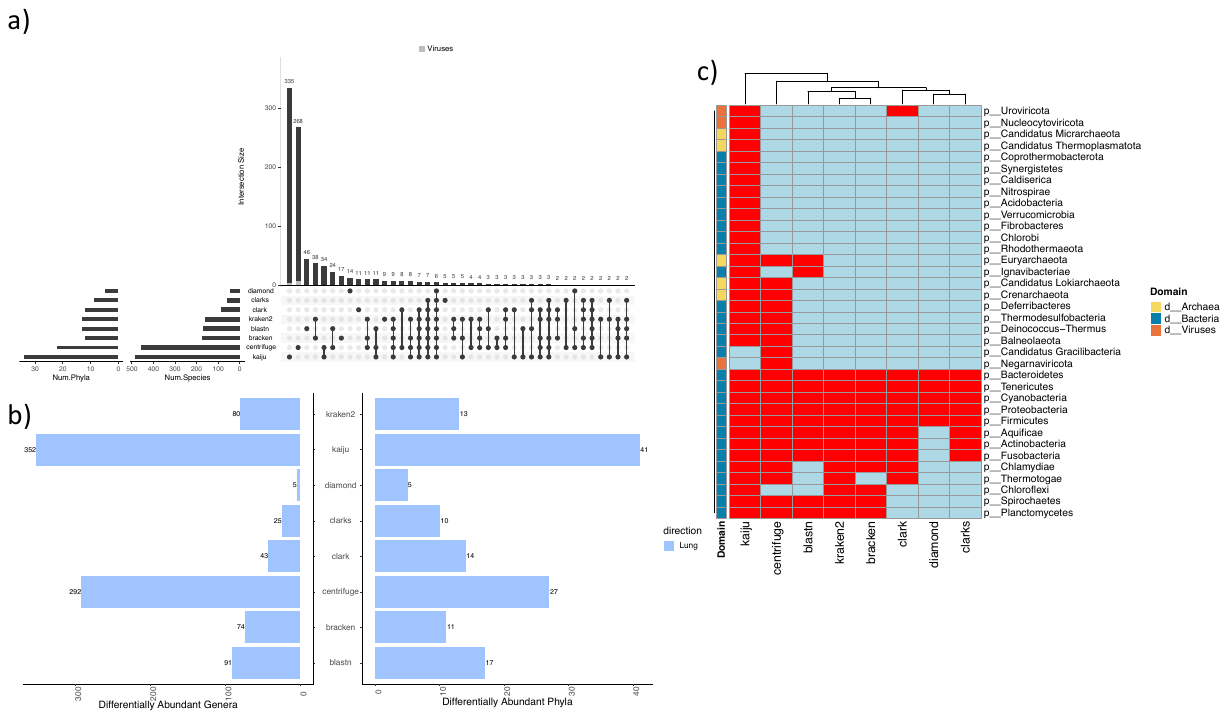

**Figure S4.** DA microbial taxa identified from comparing the microbial profiles of all rats’ kidney and spleen tissues. The number of DA species and phylum identified using different software’s profile is shown at the bar plot directly left to the software names in a). The intersection between DA taxa identified by different software at the species level is shown at the barplot at the top of a), where the number of DA virus taxa were colored in gray. Dotplot at the bottom shown the combinations of intersections between software. Number of DA taxa at phylum and genus level is shown in b). The DA phylum taxa identified by each software is shown in c), where red (
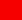
)indicate the phylum taxa at each row is reported as differentially abundant by the classification of the software in every column, and blue (
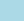
) is not reported as DA taxa by the software.

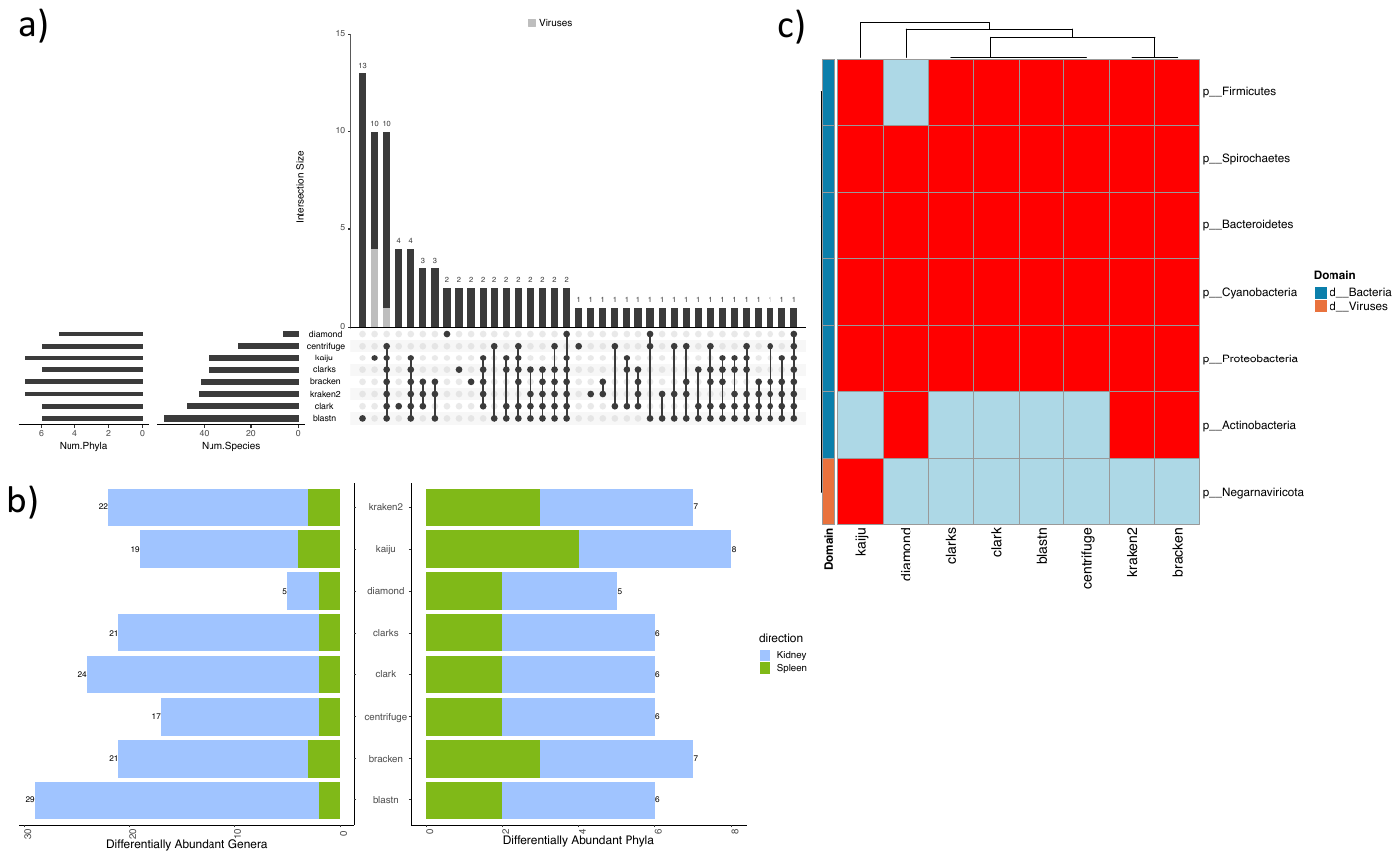
